# Genetic regulation of the human plasma proteome in 54,306 UK Biobank participants

**DOI:** 10.1101/2022.06.17.496443

**Authors:** Benjamin B. Sun, Joshua Chiou, Matthew Traylor, Christian Benner, Yi-Hsiang Hsu, Tom G. Richardson, Praveen Surendran, Anubha Mahajan, Chloe Robins, Steven G. Vasquez-Grinnell, Liping Hou, Erika M. Kvikstad, Oliver S. Burren, Madeleine Cule, Jonathan Davitte, Kyle L. Ferber, Christopher E. Gillies, Åsa K. Hedman, Sile Hu, Tinchi Lin, Rajesh Mikkilineni, Rion K. Pendergrass, Corran Pickering, Bram Prins, Anil Raj, Jamie Robinson, Anurag Sethi, Lucas D. Ward, Samantha Welsh, Carissa M. Willis, Alnylam Human Genetics, AstraZeneca Genomics Initiative, Biogen Biobank Team, Bristol Myers Squibb, Genentech Human Genetics, GlaxoSmithKline Genomic Sciences, Pfizer Integrative Biology, Population Analytics of Janssen Data Sciences, Regeneron Genetics Center, Lucy Burkitt-Gray, Mary Helen Black, Eric B. Fauman, Joanna M. M. Howson, Hyun Min Kang, Mark I. McCarthy, Eugene Melamud, Paul Nioi, Slavé Petrovski, Robert A. Scott, Erin N. Smith, Sándor Szalma, Dawn M. Waterworth, Lyndon J. Mitnaul, Joseph D. Szustakowski, Bradford W. Gibson, Melissa R. Miller, Christopher D. Whelan

## Abstract

The UK Biobank Pharma Proteomics Project (UKB-PPP) is a collaboration between the UK Biobank (UKB) and thirteen biopharmaceutical companies characterising the plasma proteomic profiles of 54,306 UKB participants. Here, we describe results from the first phase of UKB-PPP, including protein quantitative trait loci (pQTL) mapping of 1,463 proteins that identifies 10,248 primary genetic associations, of which 85% are newly discovered. We also identify independent secondary associations in 92% of *cis* and 29% of *trans* loci, expanding the catalogue of genetic instruments for downstream analyses. The study provides an updated characterisation of the genetic architecture of the plasma proteome, leveraging population-scale proteomics to provide novel, extensive insights into *trans* pQTLs across multiple biological domains. We highlight genetic influences on ligand-receptor interactions and pathway perturbations across a diverse collection of cytokines and complement proteins, and illustrate long-range epistatic effects of *ABO* blood group and *FUT2* secretor status on proteins with gastrointestinal tissue-enriched expression. We demonstrate the utility of these data for drug target discovery by extending the genetic proxied effect of PCSK9 levels on lipid concentrations, cardio- and cerebro-vascular diseases, and additionally disentangle specific genes and proteins perturbed at COVID-19 susceptibility loci. This public-private partnership provides the scientific community with an open-access proteomics resource of unprecedented breadth and depth to help elucidate biological mechanisms underlying genetic discoveries and accelerate the development of novel biomarkers and therapeutics.

## Main

Genetic studies of human populations are increasingly used as research tools for drug discovery and development. These studies can facilitate the identification and validation of therapeutic targets^1, 2^, help predict long-term consequences of pharmacological intervention^3^, improve patient stratification for clinical trials^4^, and repurpose existing drugs^5^. Several precompetitive biopharmaceutical consortia have recently invested in population biobanks to accelerate genetics-guided drug discovery, enhancing massive-scale phenotype-to-genotype studies such as the UK Biobank (UKB)^6, 7^ with comprehensive multi-omics profiling of biological samples^8–10^.

Ongoing private-public investments in biobank-based genetics are supported, in part, by a series of systematic analyses of historical drug development pipelines, all indicating that drugs developed with supporting evidence from human genetics are at least twice as likely to be approved^11, 12^. Recent advances, such as the genetics-guided repurposing of drugs targeting *IFNAR2* and *ACE2* for early treatment of COVID-19^13^ and the identification of protective, protein-truncating variants implicating *GPR75* as a therapeutic target for obesity^14^, further highlight the promise of these investments. Nonetheless, human genetics remains an imprecise instrument for biopharmaceutical research and development, as genome-wide association studies (GWAS) frequently implicate genetic variants without clear causal genes mediating their impact(s)^15^ or map to genes implicating putative drug targets with poorly understood biology or unclear mechanisms of modulation^1^.

Combining human genetics with high-throughput proteomics could help bridge the gap between the human genome and human diseases^16^. Circulating proteins can provide insights into the current state of human health^17^ and partially capture the influences of lifestyle and environment on disease pathogenesis^18^. Measuring thousands of proteins at population scale could improve genetic loss-of-function predictions^19^, help discover novel clinical biomarkers for improved patient stratification^16^, and improve fine-mapping of causal genes linked to complex diseases^2, 15^.

To date, most large-scale investigations have characterized genetic influences on blood plasma protein abundances using high-throughput aptamer^20–24^- or antibody-based^22, 25, 26^ assays. These studies have identified upwards of 18,000 associations between sequence variants and plasma protein concentrations (protein quantitative trait loci, pQTLs), using samples typically sourced from databases with proprietary subject-level access. The open-access framework^27^, deep phenotypic characterization^6^, and long-term development^8, 9, 28^ of population studies like UKB offers a unique opportunity to expand proteo-genomics to massive scale, broaden research use of high-throughput proteomic data, build more extensive pQTL databases, and accelerate the discovery of biomarkers, diagnostics and medicines. To fulfil these aims, we formed the UK Biobank Pharma Proteomics Project (UKB-PPP) - a precompetitive consortium of 13 biopharmaceutical companies funding the generation of multiplex proteomic data using blood plasma samples from UKB. Here, we describe the measurement, processing, and downstream genetic analysis of 1,472 plasma analytes measured across 54,306 UKB participants using the antibody-based Proximity Extension Assay^29^.

## Results

### Overview of UKB-PPP characteristics

We conducted proteomic profiling on blood plasma samples collected from 54,306 UKB participants using the Olink Explore 1536 platform, measuring 1,472 protein analytes, capturing 1,463 unique proteins (**Figure 1a, Supplementary Information, Extended Data Figure 1**). This included a randomised subset of 46,673 UKB participants at baseline visit (“randomised baseline”), 6,385 individuals at baseline selected by the UKB-PPP consortium members (“consortium-selected”) and 1,268 individuals who participated in the COVID-19 repeat imaging study at multiple visits **(Figure 1a, Methods).**

**Figure 1.**
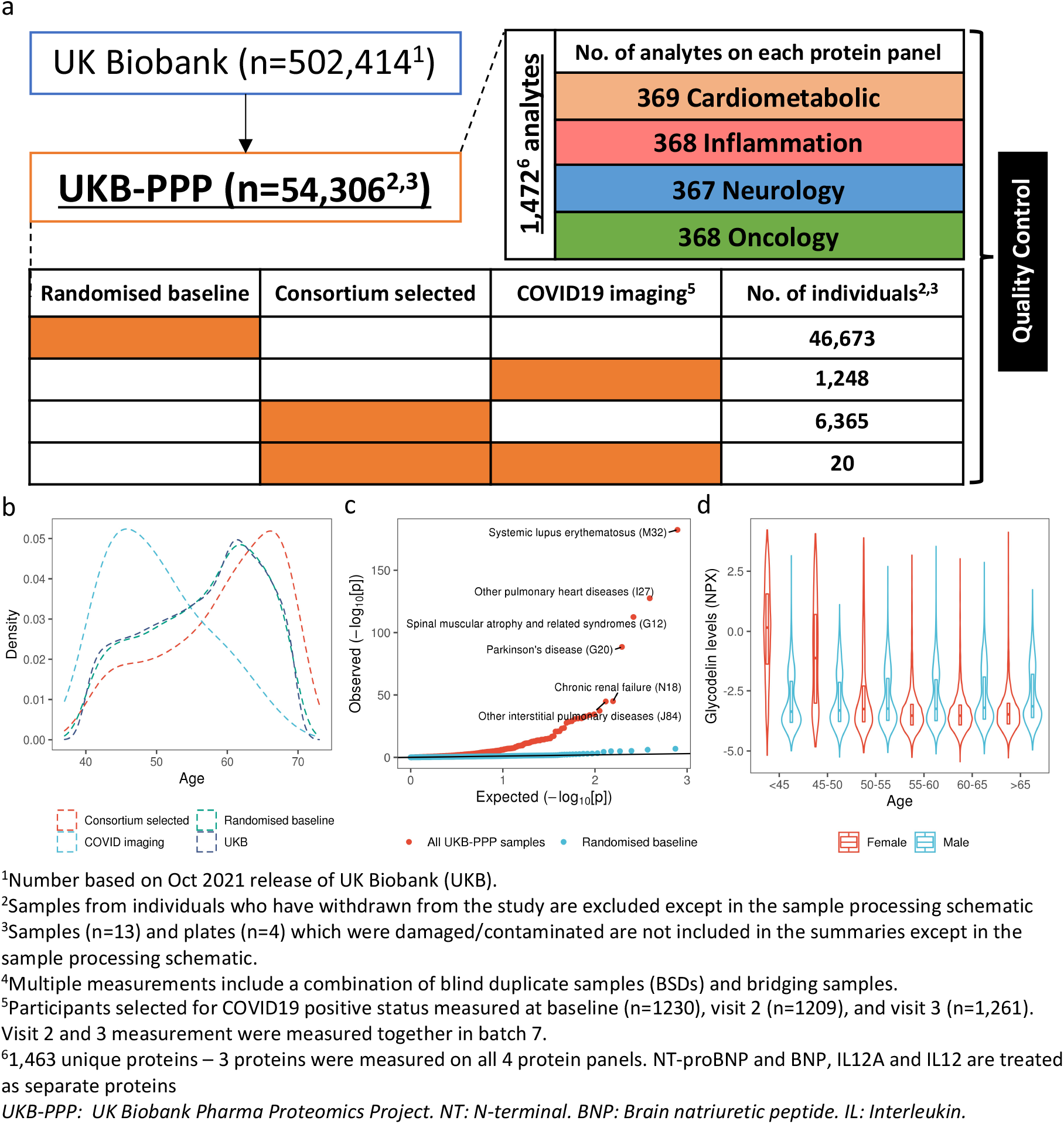
Overview of UKB-PPP. (a) Sample set-up and protein measurements. (b) Age distribution between different sub-cohorts. (c) Q-Q plot of enrichment *p*-values of UKB compared against all UKB-PPP samples and UKB-PPP randomised baseline samples. (d) Violin-plot of glycodelin (PAEP) levels by age bins and sex.

The randomised baseline participants were highly representative of the overall UKB population for various demographic characteristics (**Supplementary Table 1**). Compared to the overall UKB participants, the consortium-selected participants were on average older (by 2.5 years, *p*=5.0×10^-117^), had lower proportion of women (by 3.2%, *p*=4.1×10^-7^), and higher body mass index (BMI, by 2.6 kg/m^2^, *p=*1.3×10^-16^), different smoking prevalence (*p*=2.1×10^-6^) and composition of self-reported ethnic background (UKB data field 21000) (*p*=3.8×10^-296^), with a higher proportion of non-white ethnicities (12% vs 6%) (**Figure 1b**, **Supplementary Table 1**). The COVID-19 imaging participants had a younger age distribution (difference in means of 6.3 years, *p*=1.2×10^-162^), lower body mass index (BMI, by 1.1 kg/m^2^, *p=*1.7×10^-20^) and smoking prevalence (*p=*2.1×10^-9^), but were comparable to the overall UKB participants in sex, ethnic background, and blood group (**Supplementary Table 1**).

Compared to the full UKB cohort, UKB-PPP participants were enriched for 122 diseases, spanning multiple systems, at a Bonferroni-corrected threshold of *p<*6.7×10^-5^ (0.05/746 diseases), with no significant depletion in the diseases tested after multiple comparison adjustment (**Supplementary Table 2, Figure 1c**). This enrichment was largely driven by the inclusion of consortium selected and COVID-19 imaging participants (**Methods)** as the enrichments were mostly attenuated when considering only the randomised baseline samples (**Figure 1c);** four diseases remained modestly enriched (1.08-1.09x) and two became depleted (0.48-0.49x) in the randomised baseline samples alone (**Supplementary Table 2)**.

### Proteomic data processing and quality control

Detailed information on the Olink assay, study-wide protein measurement, processing and quality control (QC) details are provided in **Supplementary Information** and outlined in **Extended Data Figure 1** and **Figure 1a.** A total of 1,463 unique proteins were measured across four protein panels (Cardiometabolic, Inflammation, Neurology and Oncology, **Figure 1a and Extended Data Figure 1**), with 3 proteins (CXCL8, IL6, TNF) captured across all four protein panels (total=1,472 protein analytes, **Supplementary Table 3**). Globally, we did not observe batch effects, plate effects or abnormalities in protein coefficients of variation (CVs) (**Supplementary Information)**. Protein CVs, representing intra-individual variability across duplicate samples, ranged from 2.4% to 25%, with a median of 6.3% (**Supplementary Table 3, Supplementary Information**). We observed reasonably strong correlations between measurements across different panels for each of the 3 proteins measured on all four protein panels (**Extended Data Figure 2a**), with mean correlations of *r*=0.96 for CXCL8 (range: 0.95-0.98), *r*=0.92 for IL6 (range: 0.88-0.95) and *r*=0.81 for TNF (range: 0.79-0.84). We also found strong correlation (*r*=0.85) for Cystatin C independently measured using the immuno-turbidimetric approach in UKB.

### Biological associations with age, sex and BMI

In total, we found 1,126, 1,180 and 1,322 associations between protein levels and age, sex and BMI (as covariates in the same model, **Methods**) respectively at a Bonferroni-corrected threshold of *p*<3.4×10^-5^ (**Extended Data Figure 3a, Supplementary Table 4**). Many of the observed associations of protein levels with age, sex and BMI are either well-established or repeatedly reported in prior studies^20, 30–34^ – such as those between age and levels of GDF15, CHRDL1, EDA2R; sex and leptin, prostasin and CGA; and BMI and leptin, IGFBP1 and IGFBP2 (**Extended Data Figure 3a, Supplementary Table 4**). Comparing association results between overlapping proteins measured using the aptamer-based SomaScan assay in the INTERVAL study^20^, we found significant correlations in relative effect sizes for age (*r*=0.45, *p=*5.3×10^-37^), sex (*r*=0.65, *p=*1.8×10^-86^) and BMI (*r*=0.67, *p=*4.4×10^-94^) (**Extended Data Figure 3b).**

We also explored interaction effects between age, sex and BMI on protein levels in the same model. In total, we found 34 proteins levels with evidence of significant interactions (*p*<3.4×10^-5^) between age, sex and BMI; 1,149 between age and sex; 463 between sex and BMI; and 531 between age and BMI (**Supplementary Table 5**). For example, we found the strongest interaction between age and sex for glycodelin, also known as progesterone-associated endometrial protein (PAEP, *p=*2.8×10^-1445^). Glycodelin is a glycoprotein expressed in mammary glands and endometrial tissues^35^. Levels of glycodelin decreased with age for females only, particularly before the age of menopause (∼50 years), whilst for males, levels steadily increased with age (**Figure 1d).** After 55 years of age, levels of glycodelin slowly increased in females at a similar rate to males. These effects are consistent with the role of glycodelin in female reproductive tissues and their associated changes in hormone levels (such as progesterone) around menopause^35^, demonstrating that the proteomic assay used in this cohort can capture physiological effects.

### Discovery of pQTLs

Discovery pQTL analyses were performed in European ancestry participants from the randomised baseline cohort (n=35,571), which was broadly representative of the full UKB cohort, with the remaining samples (n=18,181) used as a replication cohort (**Figure 1b-c, Supplementary Tables 1-2**, **Methods**). We performed pQTL mapping of up to ∼22.6 million imputed autosomal variants for 1,463 proteins post-QC, of which 1,425 proteins are encoded by genes on autosomes. We identified 10,248 significant primary associations across 2,928 independent genetic regions at a multiple-corrected threshold of *p<*3.4×10^-11^ (**Figure 2a, Supplementary Table 6**). At a less stringent, single-phenotype genome-wide significance threshold of *p*<5×10^-8^, we found 9,150 additional associations for a total of 1,421 proteins. We base the ensuing results on associations that remained significant after adjustment for multiple testing, unless otherwise indicated. 1,377 of the 1,463 proteins tested (93.7%) had at least one pQTL at *p<*3.4×10^-11^, with 82% of proteins tested (1,162 of 1,425 proteins encoded by genes on autosomes) having a *cis* association (within 1Mb from the gene encoding the protein). We found a significant negative relationship between the number of pQTLs and the proportion of samples that were below limits of detection (LOD) for the proteins of interest (Spearman’s ρ=- 0.47, *p*= 2.7×10^-82^, **Extended Data Figure 4a**), where 67% of proteins without a pQTL (*c.f.* 3.7% of proteins with pQTL(s)) have more than 50% of samples below LOD (**Extended Data Figure 4b**). We observed, on average, a median of 6 primary associations (5^th^-95^th^ quantiles: 1-19) per protein, with 56 proteins (3.8%) having ≥20 associations (**Figure 2b top)**. Genomic inflation was well-controlled, with median λ_GC_=1.04 (standard deviation=0.018). The general inverse trend between effect size magnitudes and MAF remained for both *cis* and *trans* associations, with *trans* associations showing smaller magnitudes of effect sizes than *cis* associations (**Figure 2c).** Approximately 5.6% (570/10,248) and 1.5% (155/10,248) of the primary associations had MAF<1% and 0.5% respectively.

**Figure 2.**
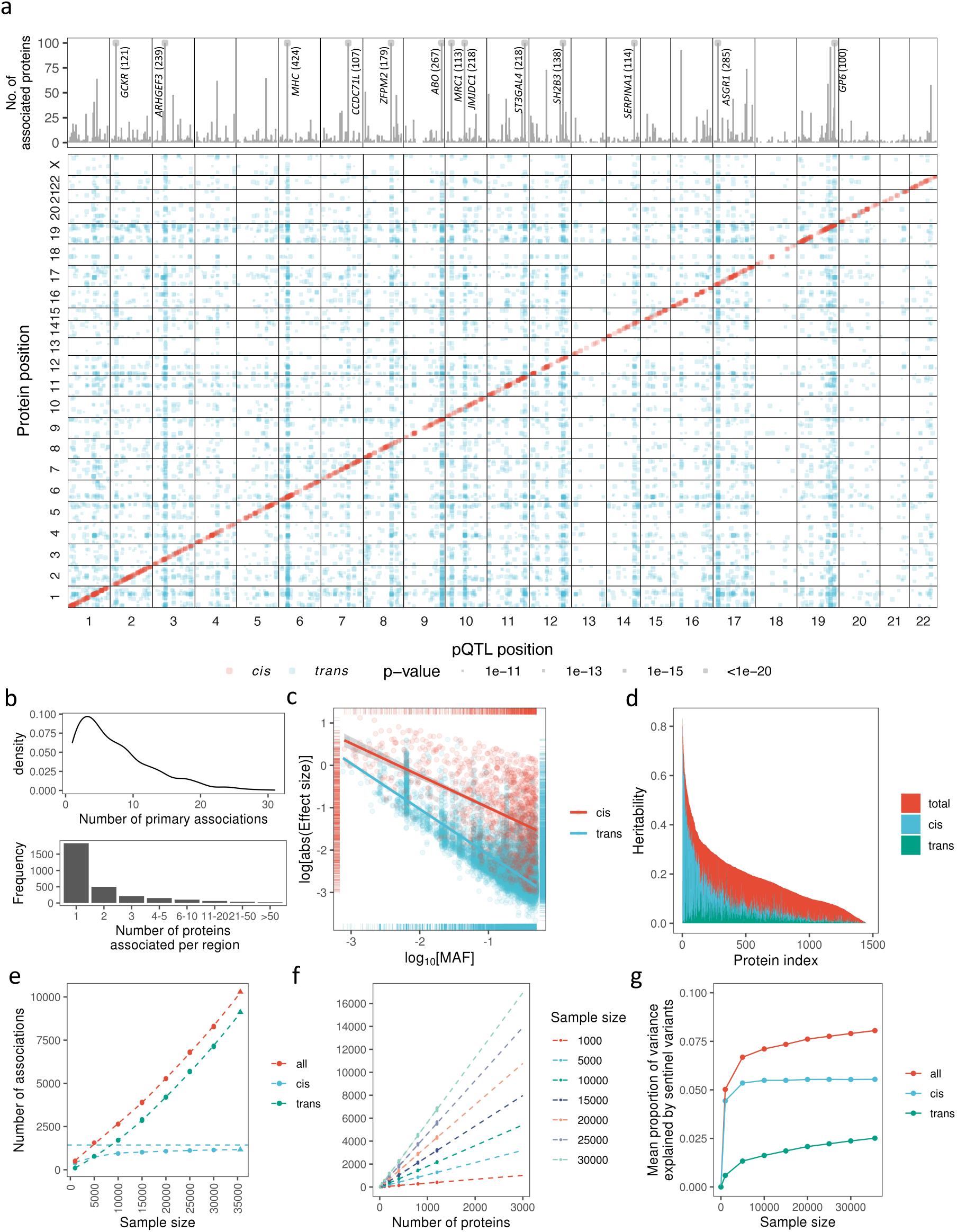
Genetic architecture of pQTLs. (a) Summary of pQTLs across the genome. Lower panel: genomic locations of pQTLs against the locations of the gene encoding the protein target. *Cis* pQTLs (red), *trans* (blue). Upper panel: number of associated protein targets for each genomic region (axis capped at 100, regions with ≥100 number of associated proteins labelled, with number in parenthesis). (b) Number of primary pQTLs per protein (top) and number of associated proteins per genomic region (bottom). (c) Log absolute effect size against log(MAF) by *cis* and *trans* associations. (d) Distribution of heritability and contributions from primary *cis* and *trans* pQTLs. (e-f) Number of primary associations against sample size (e) and number of proteins assayed (f). (g) Mean proportion of variance explained by primary pQTLs aginst sample size.

1,163 of the 10,248 primary associations were in *cis* and 9,085 were in *trans* (>1Mb from the gene encoding the protein). 59%, 95% and 97% of the *cis* associations were within the gene, 50Kb and 100Kb from the gene start site respectively. We found no systematic enrichment of *trans* pQTLs occurring on the same chromosomes as the protein tested after accounting for chromosome lengths (Fisher’s test *p*=0.89). All but two *trans* pQTLs on the same chromosome as the gene encoding the protein were >2Mb away from the corresponding gene (93% were >5Mb, 81% were >10Mb away).

63% (1,835/2,928) of the independent genetic loci were associated with a single protein, whilst 10% were associated with ≥5 proteins (pleiotropic region), and 13 loci were extremely pleotropic, associated with ≥100 proteins (**Figure 2a)**. These included well-established pleiotropic loci such as *MHC, ABO, ZFPM2, ARHGEF3, GCKR, SERPINA1, SH2B3* and *ASGR1,* all of which have previously been identified in large multiplex pQTL studies^20, 22–24^.

From the annotations of the primary pQTLs (**Extended Data Figure 5)**, we identified 25 *cis* pQTLs annotated as potential high-impact variants (e.g., frameshift, stop gained, start lost, splice acceptor, splice donor, nonsense variants) (**Supplementary Table 7)**. Among them, 10 of the primary *cis* pQTLs variants code for start codon lost/stop codon gained, of which 9 have minor alleles leading to decreased corresponding protein levels (**Supplementary Table 7**). 18 *trans* pQTLs SNPs were also annotated as potential high-impact. The majority of pQTLs identified in this study were located at non-coding regions. These non-coding pQTLs were enriched in regulatory regions, including SNPs located at promoters, enhancers, transcription factor binding sites, CTCF binding sites, and open chromatin regions (hypergeometric test *p*=3.1×10^-6^**; Supplementary Table 8**). Of the *cis* pQTLs, 23% (273) were protein-altering variants, or in LD (r^2^>0.8) with protein-altering variants (**Supplementary Table 9)**. Overall, at 49% (575) of primary *cis* associations, the index variant was in at least weak LD (r^2^>0.01) with a protein-altering variant.

### Replication of pQTLs

96.6% (9,901/10,248) of all primary associations from the discovery cohort (99.9% [1,162/1,163] *cis* and 96.2% [8,739/9,085] *trans* associations) were also nominally significant (*p*<0.05) and directionally concordant in the replication set of 18,181 participants in UKB-PPP (**Methods, Supplementary Table 6**). After adjusting for the number of associated unique genomic regions (*p<*8.7×10^-6^), 95.7% (1,113) of *cis* and 60.3% (5,480) of *trans* associations remained significant and directionally concordant in the replication cohort, inline with previous large-scale studies^20, 22–24^. Effect sizes were well-aligned between discovery and replication sets (r=0.99, *p<*10^-300^, **Extended Data Figure 6a**). Additionally, we observed good concordance of genetic associations between the three proteins measured across all four protein panels (CXCL8, IL6, TNF; **Extended Data Figure 2b),** reflecting their phenotypic correlations (**Extended Data Figure 2a).** The sentinel primary associations for these proteins were at least nominally GWAS significant across all other protein panels, suggesting good reproducibility of the same protein targets.

#### Identification of novel pQTLs

We cross referenced pQTLs identified in this study with multiple previously published pQTL results (**Supplementary Information, Methods),** finding that 85% of the primary associations from the discovery cohort (9,098/10,248) had not been identified by a prior pQTL study (**Supplementary Table 10**). A larger percentage of *trans* pQTLs were novel (91%; 9,309/10,248) than *cis* pQTLs (48%; 562/1,163).

### SNP-based heritability and variance explained by pQTLs

We estimated SNP-based heritability as a sum of contributions from significant lead pQTLs (pQTL component) and the remaining SNPs across the genome (excluding the pQTL region), which assumes a polygenic model (polygenic component) using the approach described in ^36^ (**Supplementary Table 11, Methods)**. The mean total SNP-based heritability was 0.18 (5-95^th^ quantiles: 0.02-0.44) (**Figure 1d**). On average, the *cis* primary pQTLs accounted for 19% of the overall heritability whilst the *trans* pQTLs accounted for 12% (**Figure 2d, Extended Data Figure 6b**). We found a significant correlation between the lead pQTL component and the polygenic component (Spearman’s ρ=0.52, *p*=4.7×10^-102^, **Extended Data Figure 6c**), with stronger correlations between polygenic component and *trans* pQTL (ρ=0.62, *p*= 1.6×10^-155^) component compared to *cis* (ρ=0.38, *p*= 3.5×10^-53^).

### Identification and fine mapping of independent signals

We identified 20,540 conditionally independent signals and performed fine-mapping using SuSiE (**Supplementary Table 12**). 92% (1,069/1,163) of *cis* regions contained more than one signal (mean 6.0 signals per *cis* region) (**Extended Data Figure 7**). For 11 proteins, there were 20 or more signals in the *cis* region, including CLUL1, KIR3DL1, and TPSAB1, which had 34, 26, and 23 distinct signals respectively. By comparison, only 29% (2,658/9,133) of *trans* regions contained more than one signal (mean 1.5 signals per *trans* region). Joint tagging between two or more causal variants by another non-causal variant can boost the significance of the non-causal variant in the marginal association^37–39^. We observed evidence for boosting at 3.3% (340) of tested associations, where the sentinel variant from the marginal analysis was not identified in any of the credible sets from the conditional analysis. Strong primary signals can mask the effect of independent signals in the same region, attenuating their significance in the marginal association^40^. We observed evidence for masking at 5.6% (1,142) of independent signals that were either not significant in the marginal analysis (*p>*0.05) or had opposite conditional effect directions compared to their marginal effect. Long-range regions such as the extended MHC locus have largely been ignored in large-scale genetic studies due to complicated LD structure. We observed 1,011 signals for 435 proteins mapping to the MHC locus, 139 of which were *cis* signals for 18 proteins. Together, these results underscore the importance of modelling all variants within an associated region for accurate signal identification.

We used fine-mapping to narrow down credible sets of causal variants for each independent pQTL signal (**Supplementary Table 13**). The 95% credible sets contained an average of 22.7 variants, and for 5,672 signals, we were able to determine the likely-causal variant. Credible sets for *cis* signals tended to be better resolved than those of *trans* signals (mean 95% credible set variants *cis*: 9.6; *trans*: 29.4), and were more likely to be fine-mapped to causality (signals with single variant in 95% credible set *cis*: 43%; *trans*: 20%).

### *Trans* associations highlight biological pathways and protein-protein interactions

#### Biological enrichment for proteins with multiple trans associations

For *trans* pQTLs associated with multiple independent regions (≥5) across the genome, we performed gene-set enrichment analyses by Ingenuity Pathway Analysis (IPA) to identify enrichment of biological functions relevant to cell-to-cell signalling, cellular development, development and process. We found enriched pathways for 201 proteins, including numerous enriched pathways in cellular activation, survival and signalling relevant to immune cells (**Supplementary Table 14**). For example, “activation of lymphocytes via IL8-signaling” was found to be enriched in *trans* pQTLs of CR2 protein. SNPs mapped to the nearest genes *TNFSF13B, EGFR, PAK2, HLA-DRB1, CR2, TNFRSF13B, RUNX1, ST6GAL1, PAX5* and *FOXO1* were associated with CR2 protein expression; these genes were also enriched in the IL8-signaling pathway that activates lymphocytes. In addition, we found enrichment in organismal injury mechanisms such as fibrosis (*trans* pQTLs associated with NCR1 and SMPD1) as well as in lipid metabolism, such as synthesis of triacylglycerol (*trans* pQTLs associated with SMPD1 and NAAA).

#### Protein interactions involving genes at trans loci and target protein

*Trans* associations may reflect protein interactions between the protein products of genes at the *trans* locus and the target protein (**Figure 3a)**. Additionally, genes at/near *trans* loci may operate within the same pathway as the target protein and modulate target protein levels (**Figure 3a).**

**Figure 3.**
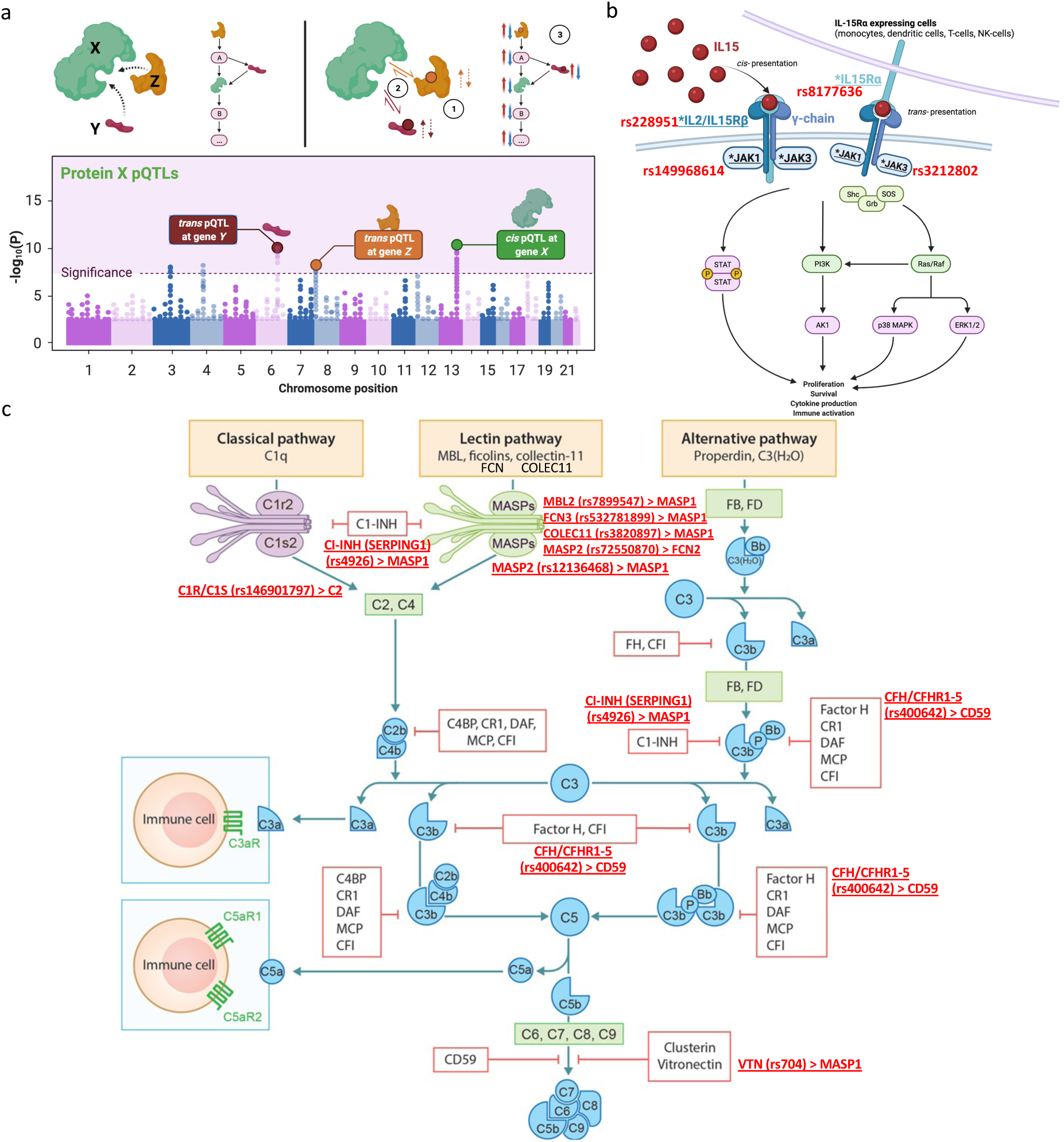
Examples of pathway networks highlighted by *trans* pQTLs. (a) Schematic of how *trans* pQTLs function as part of the same protein-protein interaction or pathway as the protein tested (protein X). Top left: proteins involved may be directly interacting or indirectly involved as part of the same pathway. Bottom: *trans* pQTLs found for corresponding genes in *trans* (in addition to potentially other signals and *cis* associations regulating protein X). Top right: some of the mechanisms by which the *trans* pQTLs may regulate the target protein (protein X) including: (1) regulating the levels of the binding partners (Y, Z) which in turn affects protein X levels, (2) altering the interaction between Y/Z with X, (3) Modulating components of the pathway in which Y/Z may be upstream/downstream of protein X. Figure created with BioRender.com including adaptations from “The Principle of a Genome-wide Association Study” (b) IL15-sginalling pathway. Components with * and underline indicate genes with *trans* pQTLs for IL15 (primary association SNP in red). Figure created with BioRender.com including adaptations from “Thrombopoietin Receptor Signaling”. (c) Complement pathway. *Trans* pQTL and associated protein in red. Figure adapted from Giang et al, Front Immunol (2018)^90^.

We used the Human Integrated Protein-Protein Interaction Reference (HIPPIE)^41^ to test if *trans* pQTL loci contained at least one gene that encoded for proteins interacting with the target protein tested. Overall, we found an interacting partner at *trans* loci for 593 proteins (**Supplementary Table 15**), including multiple receptor-ligand relationships. We found different gene products at the same pleiotropic *trans* loci interacting with different proteins with associations in those regions, which may explain certain pleiotropic effects. For 810 *trans* associations, we found a single, specific interacting protein candidate (**Supplementary Table 15)**. We also found 13 cases where the protein tested interacted with a protein in one of its *trans* loci and vice versa, indicating established coupled interactions. For example, in the ADAMTS13-vWF axis, which plays a key role in thrombosis, we found ADAMTS13 levels to be associated with a *trans* pQTL (rs112814955) at the gene encoding von Willebrand factor (*VWF*) – the substrate of the ADAMTS13 enzyme. Reciprocally, we found the *trans* pQTL for vWF (rs505922) in the *ABO* region to be 141Kb upstream of *ADAMTS13*. Other reciprocating examples included BAG3-HSPB6, PLAU-PLAUR (UROK-UPAR), TNFB-TNR1A-TNR3, GAS6-AXL, MUC16-MSLN and ITGP2-ITGAM, which are well-established protein complexes, receptor-ligand pairings, and membrane complexes. Two less well studied interactions included TNXB-APP and COL18A1-C1QTNF1, underlining potential coupled pathways for further investigation.

Notably, in addition to the HSPB6 *trans* pQTL at the *BAG3* locus (rs2234962; Cys151Arg), we found *trans* associations for both proBNP (NPPB) and NT-proBNP. BAG3 functions through BAG3-HSP70-HSPB complexes, which play an important role in heart failure and cardiomyopathies^42^, including the same *BAG3* signal (rs2234962) in previous GWAS of cardiomyopathies^43, 44^. ProBNP and NT-proBNP are established biomarkers of heart failure and cardiac damage^45^. The rs2234962 pQTL is an independent secondary *cis* pQTL for BAG3 levels from the primary *cis* pQTL (rs35434411, **Supplementary Table 12**), for which we did not find significant evidence of association with ProBNP (*p=*0.44) and NT-proBNP (*p*=0.058) levels. Taken together, these results provide additional evidence of the *BAG3* rs2234962 missense variant affecting BAG3-HSPB6 complexing, emphasizing the relevance of BAG3 to downstream blood biomarkers of heart failure and potentially cardiomyopathies.

#### Insights into cytokine and complement interactions and pathways

We found multiple instances of receptor-ligand interactions at *trans* loci for circulating cytokines and TNF superfamily proteins/receptors (**Supplementary Table 16**). In addition to *trans* pQTLs for IL15 at genes encoding its receptor components (IL15RA and IL15RB), we also found *trans* pQTLs at both *JAK1* and *JAK3,* which are proximal components of IL15 signalling (**Figure 3b);** notably, the *trans* pQTL at *JAK1* is a rare missense mutation (rs149968614, MAF=0.2%, Val651Met). Furthermore, we found that the variant rs4985556-A, which causes a premature stop gain in *IL34*, is associated with decreased levels of IL34 in *cis* (beta=-1.07, *p=*2.0×10^-1853^) and decreased CD207 (also known as langerin) - a protein marker expressed in Langerhans cells - levels in *trans* (beta=-0.08, *p=*7.4×10^-16^). Whilst IL34 and CD207 do not directly interact, this result is highly consistent with the crucial role of IL34 in development and survival of Langerhans cells^46^.

In the complement pathway, we found multiple *trans* pQTLs in genes for various constituents within the same complement pathway as the protein tested (**Figure 3c**). In particular, for protein MASP1, we found 6 of the 13 *trans* associations to lie in genes encoding other components of the complement pathway (including lectin pathway genes *MASP2, MBL2, FCN3, COLEC11,* C1-inhibitor gene *SERPING1,* and *VTN*), all of which, except *VTN,* show direct interactions with MASP1 (**Figure 3c, Supplementary Table 15)**. Notably, the *trans* pQTL at *FCN3* is a low-frequency frameshift variant (rs532781899, MAF=1.4%) leading to FCN3 deficiency^47–50^, and here, to reduced MASP1 levels (beta=-1.17, *p=*1.6×10^-328^). Similarly, we found a low frequency missense variant in *MASP2* (rs72550870, Asp120Gly, MAF=3.1%), previously linked to MASP2 deficiency^51–53^, associated with reduced FCN2 levels in this study (beta=-0.21, *p=*9.3×10^-32^). We also found C2 levels to be associated with a *trans* pQTL at *C1R/C1S* and CD59 levels with a *trans* pQTL in the *CFH-CFHR1-5* locus (**Figure 3c)**.

### Scaling of pQTL associations with increasing sample size and numbers of proteins assayed

Previous studies have performed pQTL mapping across different sample sizes and varying numbers of proteins. Here, through sub-sampling of participants and proteins, we investigated how the number of associations scaled with sample size and number of proteins assayed **(Figure 2e).** We observed an initial increase in detectable *cis* pQTLs at sample sizes below 5,000 before slowly plateauing as the number of *cis* pQTLs trended towards the number of proteins tested (1,463) – the upper bound. However, *trans* pQTLs continued to increase with larger sample sizes, without signs of plateauing at ∼54,000 participants.

Overall, the number of associations scaled linearly with the number of proteins measured (**Figure 2f)** with no obvious signs of plateauing for the current extent of proteome coverage. We found the mean proportion of variance explained by primary sentinel variants increased the most at sample sizes less than 5,000 (**Figure 2g**). Mean variance explained by *cis* associations quickly plateaued beyond samples sizes >5,000 whilst the mean variance explained by *trans* associations continued to slowly increase and drive most of the increase in mean variance explained at sample sizes >5000 (**Figure 2g**).

We also found a shift towards an increasing number of genomic regions harbouring associations with multiple proteins with larger sample sizes, indicating greater detectability of pleiotropic loci at increased study sizes (**Extended Data Figure 8a**). Furthermore, we found a slightly sublinear increase in *trans* associations with genes encoding an interacting protein with the protein tested as sample size increased **(Extended Data Figure 8b)** – suggesting further *trans* target interacting loci to be found with larger studies.

Of the four *trans* pQTLs associated with IL15 levels in the IL15 signalling pathway, associations at the *IL15RA, IL2RB, JAK1, JAK3* loci would not have been detected (*p*<3.4×10^-11^) at sample sizes below 25,000, 10,000, 20,000 and 15,000, on average, respectively. Moreover, of the 6 *trans* associations for MASP1 in the complement pathway, associations at the *MASP2, MBL2, FCN3, COLEC11, SERPING1* and *VTN* loci would not have been detected at sample sizes below 5,000, 1,000, 1,000, 1,000, 5,000, 10,000, on average, respectively. Hence, larger sample sizes would likely lead to increased discovery of *trans* pQTLs networks as opposed to isolated *trans* associations.

### Sensitivity analyses of pQTLs

We also explored, *a priori,* the impact of blood cell composition, BMI, seasonal and fasting time before blood collection on pQTL effects (**Supplementary Table 17, Extended Data Figure 9)**, discussed in more detail in **Supplementary Information.** Overall, the variables tested in the sensitivity analyses had limited impact on the majority of pQTLs.

### Co-localization with expression QTLs

Integrating pQTL results from UKB-PPP with expression quantitative trait loci (eQTL) estimates from the eQTLGen^54^ GTEx (v8)^55^, we found that 36% (507/1,425 genes available) of proteins shared a casual variant with the corresponding gene expression using the HyPrColoc method^56^ (based on a posterior probability (PP) ≥ 0.7) **(Supplementary Table 18-19)**. 11% (111/1,023 genes available) colocalized with an eQTL in whole blood (**Supplementary Table 18)** and 32% (457/1,425 genes) colocalized with eQTL(s) in at least one tissue type **(Supplementary Table 19)**. Of all targets which provided evidence of colocalization with eQTLs from the eQTLGen and GTEx consortia, 191 protein targets provided evidence of colocalization with gene expression in only one of the 49 tissues analysed (**Extended Data Figure 10a**).

Comparing the directions of effect for lead *cis* pQTL for colocalizing protein-expression combinations across tissues revealed that these were typically concordant with respect to circulating proteins and gene expression levels (**Extended Data Figure 10b)**, with 93.7% of eQTLGen and 83.6% of GTEx protein-expression combinations sharing the same direction of effect. Pervasive discordant directions of effect for molecular QTLs on gene expression and protein levels are an established phenomenon throughout the human genome, which has been postulated to be attributed to factors such as protein degradation and genetic canalization^22, 57^. Other possible explanations for discordant directions of effect include the blood-brain barrier, which may be relevant for genes such as *PARK7*, whose circulating protein shared the same direction of effect with its gene expression in 4 tissues (esophagus mucosa, heart atrial appendage, spleen and whole blood) but the opposite direction of effect in the cerebellum (**Extended Data Figure 10c**).

### Specific insights into disease, biology and potential drug targets

#### Proteomic insights into COVID-19 associated loci

The COVID-19 pandemic continues to accelerate research into the mechanisms and pathways influencing risk of COVID-19 infections and potential target candidates for drug compounds. Here we integrated pQTL data with the largest GWAS meta-analysis of reported and hospitalized COVID-19 cases conducted to date (https://www.covid19hg.org/results/r7/) using multi-trait colocalization under the HyPrColoc framework^56^.

For three of the COVID-19 hospitalization loci, we found high posterior probability of colocalization (PP>0.9) with pQTLs for proteins enriched for expression in lungs, including surfactant protein D (SFTPD), lysosome-associated membrane glycoprotein 3 (LAMP3) and mesothelin (MSLN) (**Supplementary Table 20)**. At the *MUC5B* locus, we found evidence of multi-trait colocalizations with SFTPD, LAMP3 and MSLN *trans* pQTLs, driven by the *MUC5B* promoter variant, rs35705950 (PP=1, **Figure 4a)**. Additionally, the *cis* SFTPD association colocalized with a COVID-19 hospitalization association at the *SFTPD* locus, driven by the *SFTPD* missense variant, rs721917 (PP=0.93). SFTPD has previously been causally implicated by Mendelian randomization studies for chronic obstructive pulmonary disorder^58^ and COVID-19 hospitalization^59^ risks. At the *SLC22A31* COVID-19 hospitalization locus, we also found colocalizations with another *trans* LAMP3 pQTL, driven by the *SLC22A31* missense variant, rs117169628 (PP=0.998). Apart from the pleiotropic *ABO* locus, all proteins showing evidence of pQTLs colocalizing with COVID19 hospitalization loci (PP>0.7; *SFTPD, MUC5B, ELF5, SLC22A31* and *TYK2 loci*; **Supplementary Table 20)** showed a 24-fold enrichment for their corresponding gene expression in the lungs (*p*=1.4×10^-4^).

**Figure 4.**
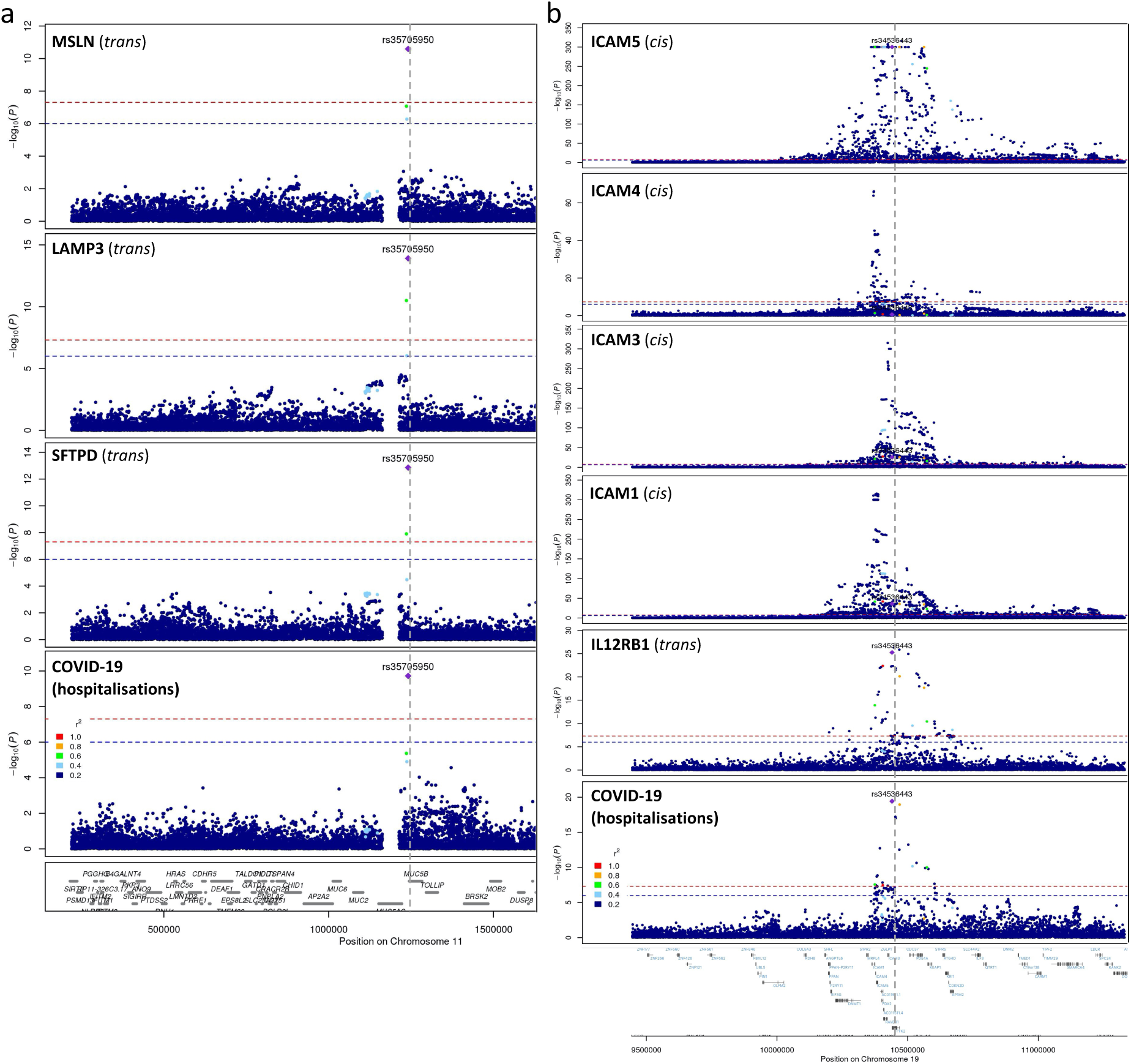
Regional association plots between COVID loci and pQTLs. (a) Regional association between COVID-19 locus at *MUC5B* and SFTPD, LAMP3, and MSLN *trans* pQTLs (b) Regional association between COVID-19 locus at *TYK2* and colocalised IL12RB1 *trans* pQTL, in addition to the *cis* pQTLs of ICAM-1,3,4 and 5 in close proximity.

In addition to colocalization at the pleiotropic *ABO* locus, we also found evidence of colocalization between the gene-dense region containing *TYK2*, ICAM-encoding genes at chromosome 19, and the interleukin-12 receptor subunit beta-1 (IL12RB1) *trans* pQTL (PP=0.95, rs34536443, *TYK2* P1104A). This pQTL is consistent with TYK2 partial loss of function caused by P1104A. No additional colocalizations were identified for the other 23 proteins with associations overlapping this locus, including ICAM-1,3,4 and 5 (**Figure 4b)**.

#### ABO blood group and FUT2 secretor epistasis effects

We observed pleiotropic associations at the *ABO* blood group and fucosyltransferase 2 (*FUT2*) loci on chromosomes 9 and 19 respectively. The FUT2 enzyme facilitates expression of ABH antigens on red cells of corresponding blood groups in mucal and gastro-intestinal (GI) secretions. Approximately 20% of white Europeans are homozygous for deletion of the *FUT2* functional secretor allele (rs601338, Trp154Ter), leading to truncation and inactivation of the enzyme and non-secretion of the blood group antigens^60^. The *FUT2* deletion has been associated with cholestatic and gastrointestinal conditions^61–63^. This led us to explore the biologically informed hypothesis that FUT2 secretor status modifies the effect of blood group antigen expression on protein levels, serving as an example of long-range gene-by-gene interaction.

We did not observe any evidence of dependencies between *ABO* blood group genotypes and FUT2 secretor status (χ^2^ *p=*0.65). At a multiple testing corrected threshold of *p<*3.4×10^-5^, 352 proteins were associated with ABO blood groups and 165 proteins were associated with secretor status (**Supplementary Table 21**). We found significant interaction between blood group and secretor status for 38 proteins. For example, CDH17, CDH1 and CGREF1 plasma levels were higher in blood group B participants compared to group A in secretors only, whilst for GALNT3, we saw the opposite effect (**Figure 5a).** We saw that the extent of differences in protein levels between secretors and non-secretors varied depended on the blood group for these proteins. We also replicated the only previous reported such interaction effect seen for alkaline phosphatase (ALP) in a Japanese cohort^64^.

**Figure 5.**
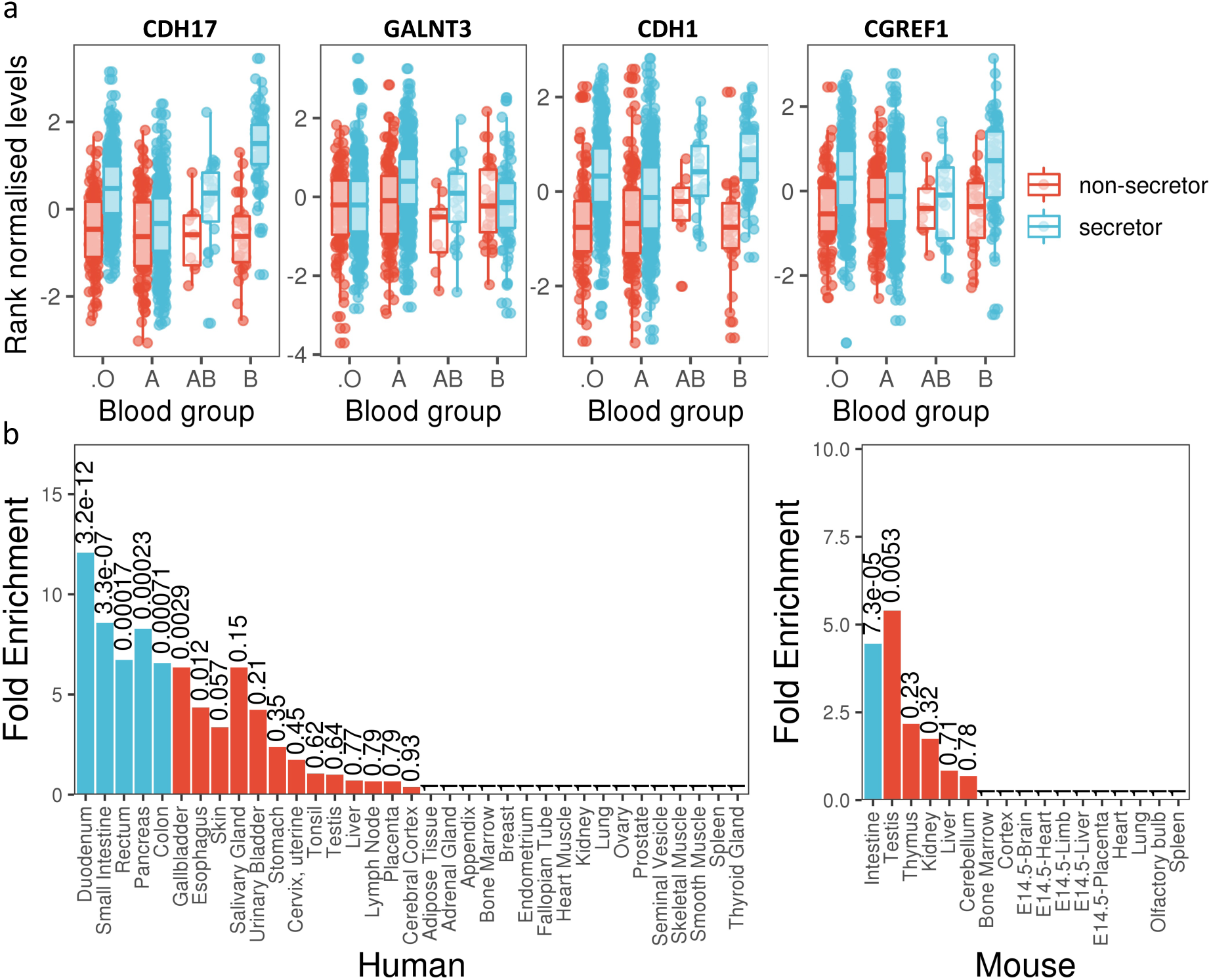
ABO blood group FUT2 secretor status interaction. (a) Boxplot of protein levels by blood group and secretor status for four proteins with most significant interaction effects. Each box plot presents the median, first and third quartiles, with upper and lower whiskers representing 1.5x inter-quartile range above and below the third and first quartiles respectively. (b) Enrichment of genes encoding proteins with significant interactions (*p*<3.4×10^-5^) for expression in various human (left) and mouse (right) tissues. Numbers above bars represent *p*-values with blue bars representing significance after multiple testing correction.

We found significant gene expression enrichments for proteins with significant interaction effects across multiple human gastrointestinal tissues^65^, including duodenum, small intestine, colon, rectum, and pancreas – consistent with the role of FUT2 in GI secretions (**Figure 5b left)**. Enrichment in the intestine was also observed in orthologous genes in a mouse tissue expression data^66^ (**Figure 5b right)**, indicating a degree of conservation between these two species.

Our results provide evidence of blood group and secretor interaction in the modulation of proteomic concentrations, which may underline susceptibility to various FUT2/ABO associated GI conditions.

#### Inflammasome pathway connections

Inflammasomes are multimeric protein complexes that mediate innate immune responses, primarily through the activation of CASP1 and subsequent cleavage, activation, and non-canonical secretion of pro-inflammatory cytokines IL-18 and IL-1β^67,68^. Rare, protein altering variants in inflammasome components are known to cause many inherited autoinflammatory conditions^69^. The causal relationship between genetic alterations in the inflammasome and autoinflammation has been clinically validated by their successful treatment with anti-IL-1β therapies^70^.

In this study, we observed multiple *trans* pQTL associations between inflammasome components and downstream effector proteins CASP1, IL-18, and IL-1β (**Supplementary Table 22)**. These associations included genes that encode inflammasome scaffolding proteins (*NLRC4, NLRP6*, and *NLRP12*); negative regulators of inflammasome activity (*VDR, CARD18*); and *GSDMD*, which enables the non-canonical secretion of IL-18 and IL-1β, and is an activator of pyroptosis **(Supplementary Table 22)**. Associations at the *NLRP12* inflammasome locus are discussed in **Supplementary Information**.

Taken together, these results indicate that - in addition to known, rare, highly penetrant, disease-causing variants – common forms of genetic variability play a more subtle, but significant, role in inflammasome-mediated innate immune responses.

#### PCSK9 pQTLs reflect pharmacological effects on cholesterol and indicated diseases

The causal effects of PCSK9 levels on LDL and total cholesterol have been well established through various orthogonal means, with several randomized clinical trials demonstrating the efficacy of PCSK9 inhibitors on cholesterol levels and cardiovascular events^71–74^. Leveraging multiple *cis* pQTLs as genetic instruments to proxy directly for the effect of PCSK9 levels, we employed Mendelian randomization to examine causal effects of PCSK9 levels on lipids (HDL, LDL and total cholesterol), cardiovascular outcomes (coronary heart disease (CHD), myocardial infarction (MI)) and ischaemic stroke (IS: large-artery (IS-LA) and small-vessel (IS-SV) subtypes) (**Methods**).

For lipids, we found significant causal effects of increased PCSK9 on increased LDL cholesterol (MR_LDL_=0.45, *p*=6.5×10^-41^) and total cholesterol (MR_TC_=0.31, *p*=4.0×10^-24^), and decreased HDL cholesterol (MR_HDL_=-0.04, *p*=0.011) (**Extended Data Figure 11**). We also found significant causal associations with increased risk of CHD (MR_log(CHD OR)_=0.24, *p*=2.2×10^-10^) and MI (MR_log(MI OR)_=0.27, *p*=9.3×10^-10^). For stroke, we found significant causal associations with increased risk of large artery ischaemic stroke subtype (MR_log(IS-LA OR)_=0.27, *p*=0.011). Whilst genetic *PCSK9* effects on LDL, total cholesterol and CHD have been found previously^36,75^, effects of PCSK9 on HDL cholesterol and large artery ischaemic stroke have not been substantiated by previous MR studies, likely due to decreased power. These findings extends the corroborated effects observed across multiple randomised clinical trials of PCSK9 inhibitors^72^.

## Discussion

High-throughput proteomic profiling of population biobanks holds the potential to accelerate our understanding of human biology and disease. Here, we present findings from one of the largest proteogenomic studies conducted to date, combining blood plasma measurements of 1,463 proteins with imputed genome-wide genotyping of 54,306 individuals in the UK Biobank. The study constructs an updated genetic atlas of the plasma proteome, identifying 10,248 primary associations with 1,377 protein levels, and provides the scientific community with an open-access, population-scale proteomics resource with individual level data and deep phenotypic integration, facilitating downstream experimentation.

We demonstrate the utility of these data for basic biological discovery using distinct examples – capturing multiple biological signalling networks, protein interactions, and long-range epistatic effects. We also underline potential use cases for drug discovery and development by validating the well-established causal relationship between PCSK9, lipid levels, cardiovascular disease and stroke, and highlight potential targets and mechanisms for COVID-19 risk. Our results expand the catalogue of genetic instruments for downstream MR and associated genomic loci for multi-trait colocalization. The availability of individual-level data should accelerate both applied and methodological studies that would not be possible with summary data. The inclusion of consortium selected samples, enriched for a range of diseases across multiple systems, also increases power for prospective proteome-disease association studies, facilitating biomarker discovery for rare conditions such as spinal muscular atrophy, where case counts are boosted by approximately five-fold compared to random sampling.

The size and breadth of this study enabled us to estimate how the genetic architecture of pQTLs scales with increasing sample size and proteome coverage, potentially guiding decisions for future proteogenomic investments. We found that the discovery of *cis* pQTLs is saturated to the number of proteins tested after ∼10,000 samples. Although *trans* association discoveries continue to increase, the heritabilities explained by *trans* loci increase at a slower rate beyond 10,000 samples. Therefore, we anticipate most gains from future, larger-scale studies to be driven by the detection of *trans* associations, rare associations and associations with proteins not previously tested. The next phase of UKB-PPP will increase the total number of plasma measurements to 2,926 unique proteins, employing the Olink Explore 3072 assay to the same individuals described in this study. We will also include 4,500 plasma samples collected approximately 10 years after initial blood draws from this randomised cohort, facilitating expanded longitudinal analyses.

Given the predominantly white European ancestral composition of UKB, the project was largely unable to capture the full genetic and phenotypic diversity of the human population. Thus, the present study and its expansion project will likely miss important insights in non-European individuals. We encourage prospective users of these data to integrate additional proteogenomic data from under-represented populations^76^, and strongly recommend that future investments in population proteomics prioritize genetic diversity in their cohort selection(s)^77^.

The study highlights the strengths of the antibody based Olink® Explore Assay for pQTL detection and downstream biological discovery. However, the Explore 1536 assay captures less than 10% of the canonical human proteome, and affinity-based platforms largely overlook protein isoforms and proteoforms generated by post-translational modifications. To address these issues, the consortium has initiated a systematic evaluation of affinity- and mass spectrometry-based assays, assessing the relative sensitivity, specificity, and scalability of the platforms, alongside the proportion of validated human proteins and proteoforms captured by each. Orthogonal validation of antibody-based proteomics using aptamer- and mass spectrometry-based assays is strongly recommended before population-scale proteomics studies expand to sample sizes of 100,000 and beyond.

Following on from the successful exome sequencing and the ongoing whole genome sequencing of UK Biobank, the Pharma Proteomics Project builds on the precompetitive industry collaboration framework in generating high-dimensional, population-scale data for the advancement of science and medicine. The wider research community will be able to leverage this open-access resource to test hypotheses crucial to the development of improved diagnostics and therapeutics for human disease.

## Methods

### UK Biobank participants

UK Biobank (UKB) is a population-based cohort of approximately 500,000 participants aged 40-69 years recruited between 2006 and 2010. Participant data include genome-wide genotyping, exome sequencing, whole-body magnetic resonance imaging, electronic health record linkage, blood and urine biomarkers, physical and anthropometric measurements. Further details are available at https://biobank.ndph.ox.ac.uk/showcase/. All participants provided informed consent. This research has been conducted using the UK Biobank Resource under approved application numbers 65851, 20361, 26041, 44257, 53639, 69804.

### UKB-PPP sample selection and processing

Details of UKB participant selection and sample handling are detailed in **Supplementary Information.**

### Proteomic measurement, processing and quality control

Details of the Olink proteomics assay, data processing and quality control are detailed in Supplementary Information.

### Genomic data processing

UKB genotyping and imputation (and quality control) were performed as described previously^6^. In addition to checking for sex mismatch, sex chromosome aneuploidy, and heterozygosity checks, imputed genetic variants were filtered for INFO>0.7, MAC>50 and chromosome positions were lifted to hg38 build using LiftOver^78^. European ancestry was defined using the Pan-UKBB definitions in UKB return dataset 2442, “pop = EUR”.

### Genetic association analyses

GWAS were performed using REGENIE v2.2.1 via a two-step procedure to account for population structure detailed in ^79^. In brief, the first step fits a whole genome regression model for individual trait predictions based on genetic data using the leave one chromosome out (LOCO) scheme. We used a set of high-quality genotyped variants: minor allele frequency (MAF)>1%, minor allele count (MAC)>100, genotyping rate >99%, Hardy-Weinberg equilibrium (HWE) test *p*>10^−15^, <10% missingness and linkage-disequilibrium (LD) pruning (1000 variant windows, 100 sliding windows and r^2^<0.8). The LOCO phenotypic predictions were used as offsets in step 2 which performs variant association analyses using standard linear regression. We limited analyses to variants with INFO>0.7 and MAC>50 to minimise spurious associations.

In the discovery cohort (n=35,571), we included participants of European ancestry from batches 1-6, and excluded the pilot batch, plates which were normalised separately, and batch 7 (COVID-19 imaging longitudinal samples and baseline samples showing increased variability and mixed with COVID-19 imaging samples). Participants not included in the discovery cohort were included in the replication cohort, which consisted of 14,706 White, 1,225 Black/Black British, 998 Asian/Asian British, 148 Chinese, 339 Mixed, 613 Other and 152 missing ethnic backgrounds based on the self-reported ethnicities in UKB (data field 21000).

For the discovery cohort, association models included the following covariates: age, age^2^, sex, age*sex, age^2^*sex, batch, UKB centre, UKB genetic array, time between blood sampling and measurement and the first 20 genetic principal components (PCs). The covariates in the replication cohort additionally included whether the participant was pre-selected, either by the UKB-PPP consortium members or as part of the COVID imaging study.

To ensure reproducibility of the analysis protocol, the same proteomic QC and analysis protocols were independently validated across two additional sites using the same initial input data on the three proteins measured across all protein panels (CXCL8, IL6, TNF).

### Definition and refinement of significant loci

We used a conservative multiple comparison-corrected threshold of *p*<3.4×10^-11^ (5×10^-8^ adjusted for 1,463 unique proteins) to define significance. We defined primary associations through clumping ±1Mb around the significant variants using PLINK^80^, excluding the HLA region (chr6:25.5-34.0Mb) which is treated as one locus due to complex and extensive LD patterns. Overlapping regions were merged into one, deeming the variant with the lowest *p*-value as the sentinel primary associated variant. To determine regions associated with multiple proteins, we iteratively, starting from the most significant association, grouped together regions associated with proteins containing the primary associations that overlapped with the significant marginal associations for all proteins (*p*<3.4×10^-11^). In cases where the primary associations contained marginal associations that overlapped across multiple groups, we grouped together these regions iteratively until convergence.

### Variant annotation

Annotation was performed by Ensembl Variant Effect Predictor (VEP), ANNOVAR (https://annovar.openbioinformatics.org/en/latest/) and WGS Annotator (WGSA, https://sites.google.com/site/jpopgen/wgsa). The gene/protein consequence was based on RefSeq and Ensembl. We reported exon and intron numbers that a variant falls in as in the canonical transcripts. For synonymous mutations, we estimated rank of genic intolerance and consequent susceptibility to disease based on the ratio of loss-of-function. For coding variants, SIFT and PolyPhen scores for changes to protein sequence were estimated. For non-coding variants, transcription factor binding site, promoters, enhancers and open chromatin regions were mapped to histone marks chip-seq, ATAC-seq and DNase-seq data from The Encyclopedia of DNA Elements Project (ENCODE, https://www.encodeproject.org) and ROADMAP Epigenomics Mapping Consortium (http://www.roadmapepigenomics.org). For intergenic variants, we mapped the 5’ and 3’ nearby protein coding genes and provided distance (from 5’ transcription starting site of a protein coding gene) to the variant. Combined Annotation Dependent Depletion score (CADD, https://cadd.gs.washington.edu) was estimated for non-coding variants. An enrichment analysis hypergeometric test was performed to estimate enrichment of the associated pQTL variants in specific consequence or regulatory genomic regions.

### Cross referencing with previously identified pQTLs

To evaluate whether the pQTLs in the discovery set were novel, we used a list of published pQTL studies (http://www.metabolomix.com/a-table-of-all-published-gwas-with-proteomics/) and the GWAS catalog to identify previously published pQTL studies. Twenty-six studies were included (**Supplementary Information**). Using a *p*-value threshold of 3.4×10^-11^, we identified the sentinel variants and associated protein(s) in the previously published studies and queried those against our discovery pQTLs. If a previously associated sentinel variant-protein pair fell within a 1Mb window of the discovery set pQTL sentinel variant for the same protein and had an r^2^≥0.8 with any significant SNPs in the region, it was considered a replication.

### Heritability analysis

We estimated the SNP-based heritability as a sum of variance explained (VE) from the primary sentinel variants for each protein at each loci (pQTL component) and the polygenic component using the genome-wide SNPs excluding the pQTL regions of each protein. The polygenic component, which mostly likely satisfies the polygenic model of small genetic contributions across the genome, was estimated using LD-score regression^81^.

### Identification and fine mapping of independent signals

We used the Sum of Single Effects model (SuSiE, version 0.12.6)^39^ to identify and fine map independent signals from subject level data. To create test regions that accounted for potential long-range LD, we performed a two-step clumping procedure using PLINK with parameters 1) “--clump-r2 0.1 --clump-kb 10000 --clump-p1 3.4e-11 --clump-p2 0.05” on the summary statistics and 2) “--clump-kb 500” on the results of the first clumping step. For each clump, we extended the coordinates of the left- and right-most variants to a minimum size of 1 Mb. We merged overlapping clumps and defined these as the test regions. For each test region, we applied SuSiE after pruning pairs of related samples (1^st^ or 2^nd^ degree relations) and regressing out the same covariates as the main analysis with parameters “min_abs_corr=0.1, L=10, max_iter=100000, refine=TRUE”. For test regions where SuSiE found the maximum number of independent signals, which was initially set at “L=10”, we incremented “L” by 1 until no additional signals were detected (up to a maximum of L=35 for the *cis*-region of CLUL1).

### Pathway enrichment and protein interactions

For pleiotropic pQTL loci and multiple associated *trans* pQTL proteins, gene-set enrichment analyses were performed by Ingenuity Pathway Analysis (IPA) to identify enrichment of biological functions relevant to cell-to-cell signaling, cellular development, development and process. Gene pathways and networks annotated based on STRING-db and KEGG pathway databases were also used for enrichment analyses. Hypergeometric tests were performed to estimate statistical significance and hierarchical clustering trees and networks summarizing overlapping terms/pathways were generated. To correct for multiple testing, the false discovery rate (FDR) was estimated. FDR < 0.01 was considered as statistical significance.

To test if *trans* pQTL loci contained at least one gene (within 1Mb of the *trans* pQTL) that encoded for proteins interacting with the tested protein, we used the curated protein interaction database: Human Integrated Protein-Protein Interaction Reference (HIPPIE)^41^ release v2.3 (http://cbdm-01.zdv.uni-mainz.de/~mschaefer/hippie/download.php).

### Sub-sampling analysis

To estimate how the number of associations scaled with sample size, we took random samples without replacement of [1,000, 5,000, 10,000, 15,000, 20,000, 25,000 and 30,000] from the discovery randomized baseline cohort, then performed the association analyses of the primary sentinel variant and examined the proteomic variance explained in the exact same manner as the main analyses described above. We also examined how associations scaled with the number of proteins measured by random sub-sampling [10, 50, 100, 200, 400, 800, 1200] proteins from the results. We also performed multiple samples (n=3) to check consistency and stability of sub-sampling results across runs.

### Sensitivity analyses

The variables for sensitivity analyses were chosen *a priori* to avoid post-hoc biases.

#### Effects of blood cell counts

We investigated the effect of blood-cell (BC) composition on the genetic association with plasma proteins through sensitivity analyses of pQTLs from the discovery analyses. The top hits from the discovery analyses were re-analysed adjusting for the following blood-cell covariates: monocyte count; basophil count; lymphocyte count; neutrophil count; eosinophil count; leukocyte count; platelet count; hematocrit percentage; hemoglobin concentration. These blood-cell covariates were selected to represent blood-cell composition due to their common clinical use. Prior to the analyses, we followed the methods in ^82^ to exclude blood-cell measures from individuals with extreme values or relevant medical conditions. Relevant medical conditions for exclusion included pregnancy at the time the complete blood count was performed, congenital or hereditary anemia, HIV, end-stage kidney disease, cirrhosis, blood cancer, bone marrow transplant, and splenectomy. Extreme measures were defined as leukocyte count >200×10^9^/L or >100×10^9^/L with 5% immature reticulocytes, hemoglobin concentration >20 g/dL, hematocrit >60%, and platelet count >1000×10^9^/L. After blood-cell measure exclusions, all individuals in the discovery cohort without blood-cell measures had each measure imputed to the mean of the cohort. Following these exclusions and QC, genetic analyses of the sentinel variant – protein associations adjusted for blood-cell covariates were performed using the same approach as the main analysis.

We further tested whether blood cell composition is partially or fully mediating variant-protein associations (Genotype -> BC measure -> Protein) for genetic associations significant within the discovery (*p<*3.4×10^-11^) and not in the sensitivity analyses (*p>*3.4×10^-11^). For each variant – protein association, we first identified the BC phenotypes that were associated with protein levels at *p*<3.4×10^-11^ within a multivariate linear regression model including blood cell phenotypes as the predictors, protein as the outcome and adjusted for all other covariates included in the discovery analysis. We then confirmed if there was an association between the genetic variant (dosage) and each of the blood cell phenotypes (Genotype -> BC) and between blood cell phenotype and the protein (BC -> Protein) prior to testing for mediation. In the final test, we compared the strength of associations, Genotype -> Protein, to that of the Genotype -> Protein in a multivariate model (Protein ∼ Dosage + BC phenotype + Discovery Covariates) to establish whether the variant – protein association is either fully (*p*>0.01) or partially (*p*<3.4×10^-11^) mediated by the blood cell phenotype.

#### Effects of BMI

We investigated the effect of BMI on the genetic association with plasma proteins through sensitivity analyses of pQTLs from the discovery analyses. The primary associations from the discovery analyses were re-analysed using the same approach as the main analysis including BMI [data field: 21001] as an additional covariate.

#### Effects of season and amount of time fasted at blood collection

To assess the effects of season and amount of time fasted at blood collection on variant associations with protein levels, we re-analysed all sentinel pQTLs identified in the main discovery analyses including season and fasting time as two additional covariates. Blood collection season (summer/autumn: June to November vs. winter/spring: December to May) was defined based on the blood collection date and time (data-field: 3166). Participant-reported fasting time was derived from data-field 74 and was standardized (Z-score transformation) prior to analysis.

### Co-localization analyses

We investigated evidence of shared genetic variation between the 1,425 circulating proteins encoded by autosomal genes and their tissue-specific gene expression using the HyPrColoc approach^56^. Analyses were conducted using variant-level priors; alignment probabilities and a posterior probability of colocalization (PP) ≥ 0.7 threshold was applied to indicate evidence of shared genetic variation. For each circulating protein in turn, we aggregated *cis* pQTL estimates around their encoding gene region (+/-500kbs) from the discovery UKB-PPP GWAS as well as *cis*-expression quantitative trait loci (eQTL) using whole blood derived findings from the eQTLGen consortium^54^ and 48 other tissue types from the GTEx consortium^55^ (v8). This included all available tissues with eQTLs in GTEx, excluding whole blood, as these data were included in the eQTLGen meta-analysis.

Next, for circulating proteins which provided evidence of colocalization in this previous analysis, we assessed whether lead *cis* pQTL influenced protein levels and gene expression in the same direction (for gene expression in tissues which provided evidence of colocalization). Lead *cis* pQTL were selected as those with the smallest *p*-value that also existed in the corresponding eQTL dataset which were not palindromic variants.

For colocalization with COVID-19 loci, the top loci reported by the COVID-19 Host Genetics consortium (https://app.covid19hg.org/variants) were updated with estimates from the R7 summary results (https://www.covid19hg.org/results/r7/) for hospitalised COVID-19 cases and reported COVID-19 infections compared to population controls.

### PCSK9 Mendelian randomization

#### Instrument selection and outcomes

Instruments to proxy for altered PCSK9 abundance were generated using variants associated in *cis* (within 1Mb of the PCSK9 gene-coding region) at genome-wide significance (*p*<5×10^-8^) to minimise pleiotropic effects. We performed LD clumping to ensure SNPs were independent (r^2^ < 0.01) by using an in-sample reference panel of 10,000 UK Biobank participants. We removed SNPs with a F-statistic less than 10 to avoid weak instrument bias.

Outcomes of interest were measurements of cholesterol, including low-density lipoprotein cholesterol (LDL-c), high-density lipoprotein cholesterol (HDL-c), triglycerides (TG) and total cholesterol (TC); coronary heart disease (CHD) and myocardial infarction (MI); ischemic stroke large artery atherosclerosis and small-vessel subtypes. Data for these outcomes were extracted from the OpenGWAS project^83, 84^. *PCSK9* pQTL effects were harmonised to be on the same effect allele. If the variant was not present in the outcome dataset, we searched for a proxy SNP (r^2^>0.8) as a replacement if available.

#### MR analysis

We performed two-sample MR on the harmonised effects to estimate the effect of genetically proxied PCSK9 abundance on genetic liability to the outcomes of interest. We estimated the effects for each individual variant using the two-term Taylor series expansion of the Wald ratio (WR) and the weighted delta inverse variance weighted (IVW) to meta-analyse the individual SNP effects to estimate the combined effect of the WRs. Results from the MR analyses were interrogated using standard sensitivity analyses. We used Steiger filtering to provide evidence of whether the estimated effect was correctly orientated from PCSK9 abundance to the outcome and not due to reverse causation.

### ABO blood group and FUT2 secretor status analysis

ABO blood group was imputed through the genetic data using three SNPs in the *ABO* gene (rs505922, rs8176719, and rs8176746) following the blood-type imputation method in UKB (https://biobank.ndph.ox.ac.uk/ukb/field.cgi?i=23165), developed from ^85–88^. FUT2 secretor status was determined by the inactivating mutation (rs601338), with genotypes GG or GA as secretors and AA as non-secretors. Interaction term between blood group (O as reference group) and secretor status was tested adjusting for the same covariates as in the main pQTL analyses for each protein separately. A multiple testing threshold of *p<*3.4×10^-5^ (0.05/1,463 proteins) for the interaction terms was used to define statistically significant interaction effects.

### Enrichment for gene expression in tissues

Tissue enrichment of associated proteins was tested using the TissueEnrich R package (v1.6.0)^89^, using the genes encoding proteins on the Olink panel as background. For enrichment in human genes, we used the RNA dataset from Human Protein Atlas^65^ using all genes that are found to be expressed within each tissue, whilst for orthologous mouse genes we used data from Shen *et al.*^66^. The enrichment *p*-value thresholds were corrected for multiple comparisons based on the number of tissues tested (n=35 in human and n=17 in mouse tissues).

## Acknowledgements

We thank the participants, contributors, and researchers of UK Biobank for making data available for this study – with special thanks to Lauren Carson, John Busby, Naomi Allen and Rory Collins for making the study possible. We are grateful to the research & development leadership teams at the thirteen participating UKB-PPP member companies (Alnylam Pharmaceuticals, Amgen, AstraZeneca, Biogen, Bristol-Myers Squibb, Calico, Genentech, Glaxo Smith Klein, Janssen Pharmaceuticals, Novo Nordisk, Pfizer, Regeneron, and Takeda) for funding the study. We thank the Legal and Business Development teams at each company for overseeing the contracting of this complex, precompetitive collaboration – with particular thanks to Erica Olson of Amgen, Andrew Walsh of GSK, and Fiona Middleton of AstraZeneca. The Biogen team is especially thankful to Helen McLaughlin for her project management support. Finally, we thank the team at Olink Proteomics (Philippa Pettingell, Klev Diamanti, Cindy Lawley, Linda Jung, Sara Ghalib, Ida Grundberg and Jon Heimer) for their consistent logistic support throughout the project – with special thanks to Evan Mills for co-championing the project and leading internal activities at Olink.

## Data availability

Full summary association data are available at [URL available on publication]. Underlying proteomics data is available under return dataset [return dataset ID and URL on publication depending on time of official publication] of UK Biobank.

## Code availability

Codes used are part of standard software and tools.

## Author contributions

Study conceptualization & project coordination: C.D.W.; study design and methodology B.B.S., C.D.W., C.B., J.C., L.H., Y.H.H., E.M.K., A.M., T.G.R., C.R., P.S., M.T., O.S.B., J.D., K.L.F., C.E.G., Å.K.H., S.H., T.L., R.M., R.K.P., B.P., J.R., N.T., S.V.G., L.W., C.M.W., M.H.B., H.M.K., E.N.S., J.D.S., B.W.G., M.R.M.; proteomic data QC: B.B.S., K.L.F., T.L.; phenotype harmonisation: B.B.S., T.L., K.L.F., L.B., S.W., C.P.; analysis: B.B.S., C.D.W., C.B., J.C., L.H., Y.H.H., E.M.K., A.M., T.G.R., C.R., P.S., M.T., O.S.B., J.D., K.L.F., C.E.G., Å.K.H., S.H., T.L., R.M., R.K.P., B.P., J.R., N.T., S.G.V., L.W., C.M.W., M.H.B., H.M.K., E.N.S., J.D.S., B.W.G., M.R.M., E.B.F.; genetic association analyses: B.B.S.; independent replication of genetic analyses: A.M., C.B., E.M.K.; Mendelian randomization: J.R.; conditional analyses: J.C.; epitope mapping analysis: A.M., C.B.; co-localization with eQTLs: T.G.R, M.T.; sensitivity analysis: A.M., C.B., C.R., P.S., L.H.; variant annotation: Y.H.H.; writing first draft of manuscript: B.B.S., C.D.W.; writing second draft of manuscript: B.B.S., A.M., M.H.B., S.V.G, Y.J., A.K.H., S.P., B.G., S.S., J.D.S., C.R., P.S., E.B.F., L.W., C.W., E.M.K., J.M.M.H., H.M.K., C.G., E.N.S., L.B.G., S.W., M.R.M., C.D.W.; all authors critically reviewed the manuscript.

## Inclusion and ethics statement

The inclusion and ethics standards have been reviewed where applicable.

