## Supplementary Information for "Genetic regulation of the human plasma proteome in 54,306 UK Biobank participants"

#### UK Biobank sample selection for UKB-PPP

Samples were selected in two temporally and algorithmically separated picking processes. For the first, initially 5,500 samples collected from participants during their baseline recruitment visit were pre-selected by the Consortium members. 44,502 further representative participant samples were selected from the UK Biobank cohort through a stratified selection against age, sex, and recruitment centre for baseline samples, optimised to reduce number of plates for the picking process. Day of week of collection, participant ethnicity and deprivation index for selected participants were confirmed as representative of cohort distributions. This created a total selection of 50,002 samples. For the second picking process a total of 7,000 samples were supplied. First, 1,020 pre-selected samples from the Consortium members were chosen. A further 3,637 samples were selected from participants attending the COVID-19 case-control imaging study. 1,270 participants formed this section of the study, who had been invited to attend UK Biobank for a repeat imaging assessment on the basis of a COVID-19 diagnosis from linked healthcare records or a positive home-based SARS-CoV-2 antibody lateral flow test provided by UK Biobank, or were invited as matched controls of these COVID-19 cases. Full inclusion and eligibility criteria are described in https://biobank.ndph.ox.ac.uk/showcase/showcase/docs/casecontrol_covidimaging.pdf.

For these 1,270 participants, samples from initial recruitment, pre-COVID and post-COVID imaging visits were included where possible. An additional 2,343 baseline samples were included in this picking process selected randomly as described for the first sample selection, optimised to include selection locations required for the COVID imaging and consortium-selected samples.

#### UKB sample handling

EDTA (9ml) vacutainers were collected and fractioned to 850µl aliquots of EDTA plasma, buffy coat and red cells as per the UK Biobank collection protocol described in [1]. Aliquots were stored in an automated -80°C sample archive. EDTA plasma from participants selected for the study were withdrawn from the automated sample archive in a quasi-randomised order[2]. Prior to processing, aliquots were stored in a -80°C freezer. Racks containing 85 aliquots were thawed (column 12 plus one additional random well left empty for controls and two spaces for blinded duplicates) and 60µl of EDTA plasma was pipetted to a PCR plate (P/N AB0800, Thermo Scientific™) using a TECAN freedomEVO with full sample tracking. Two blinded duplicates (using aliquots from the same plate) were added to each plate before plates were sealed with adhesive seals (P/N 4306311, Applied Biosystems™) and stored at -80°C. Samples were shipped on dry ice to Olink Analysis Service in Sweden for analysis accompanied by an electronic sample manifest containing sample level information.

#### Plasma profiling using the Olink technology

The Olink technology uses Proximity Extension Assay, where a matched pair of antibodies labelled with unique complimentary oligonucleotides (proximity probes) bind to the respective target protein in a sample. As a result, the probes come into close proximity and hybridize to each one, enabling DNA amplification of the protein signal, which is quantified on a next generation sequencing read-out which has been described in detail previously[3,4]. Antibodies targeting 1463 unique proteins are distributed across four 384-plex panels and each panel composed of four dilution blocks to accommodate for the different dynamic ranges of target proteins in plasma and serum. The panels focus on inflammation (INF), oncology (ONC), cardiometabolic (CAM) and neurological (NEU) proteins. A summary of all proteins is shown in **Supplementary Table 3** and a schematic of the assay approach is summarised in **Extended Data Figure 1**. The performance of each protein assays is validated based on specificity, sensitivity, dynamic range, precision, scalability detectability and endogenous interference[3].

Samples were serially diluted to 1:10, 1:100, 1:1000, transferred to four 384-well plates consisting of four abundance blocks for each of the four panels per 96 samples. In the immune-reaction, plasma samples were incubated overnight at 4°C with the proximity probes. Oligonucleotides in close proximity are extended and amplified using DNA polymerase and create a DNA sequence which is amplified by polymerase chain reaction (PCR 1), to create amplicons encoding protein assay information. All amplicons for each sample from the four abundance groups per panel are combined, resulting in one well of amplicons for each sample/panel. Four unique 96- index plates are added to every 4 sample plates, followed by a second PCR (PCR 2), to enable all samples in a plate to be combined in to one library per panel. Each library is bead purified and libraries quality controlled using a Bioanalyzer. Four sample plates from four identical panels were pooled, denatured and sequenced on individual lanes on the Novaseq600 using S4 flow cells v1.5 (35 cycles) and 384-samples and 384-assays measured. Counts of known sequences were translated in to Normalized Protein eXpression (NPX) values within Olink's MyData Cloud Software.

***Olink inbuilt assay quality control***

The Olink workflow has an inbuilt quality control system, 3 engineered internal controls that are spiked into every sample, and each abundance block. The incubation control (Inc Ctrl) is a non-human assay, green fluorescent protein (GFP) and is used during data QC. The extension control consists of two paired-oligonucleotides coupled to an antibody molecule with the DNA arms in close proximity and was used for normalization of the data. The amplification control (Amp Ctrl) consists of a synthetic double stranded DNA template, used for QC and to monitor the PCR steps in the protocol. In addition, each sample plate includes external controls in column 12. A negative control ran in triplicate is used to calculate the limit of detection (LOD) of each assay in every plate, and a plate control sample, consisting of a pooled plasma sample is run in triplicate to adjust the levels between plates. A duplicate pooled sample control is included to estimate precision within and between runs. In each of the four panels, 3 overlapping assays of IL6, IL8 (CXCL8), and TNF are included for quality control (QC) purposes to assess correlation.

Olink's internal QC assessment is performed at two levels; run QC and sample QC. For run QC, each abundance block per panel and sample plate should fulfil the mean absolute deviation (MAD) in both internal controls (Inc Ctrl and Amp Ctrl) which should not exceed 0.3 NPX, the deviation of sample QC level is allowed for up to 1/6 samples and in each panel the median of 90% assays in plate and negative controls should be in the accepted range from predefined values set during validation. The sample QC assesses all samples individually using the internal controls (Inc Ctrl and Amp Ctrl) which should be within ± 0.3 NPX from the plate median across the abundance block, in addition to the mean assay count for a sample may not be less 500 counts. Samples that don't fulfil these criteria will receive a sample warning for a given abundance block in the data set. Data from these assays should be treated with caution. Assays where the median of the negative Control triplicates deviate more than 5 SDs from predefined values set for each assay during validation will receive a QC warning in the results NPX file. Data was generated according to Olink's standard procedures. Normalized Protein eXpression (NPX) is Olink's relative quantification unit on a log-2 scale. Data generation consists of normalization of matched counts of an assay to the extension control spiked into every sample, log-2 transformation of the data, and level adjustment using the plate control.

#### NPX calculation and normalization

Samples from the study were divided into two sets: i) Set 1 – UKB; and ii) Set 2 – COVID; depending on the time point they were randomly selected from the UKB population. Samples were randomly assigned to 96-well plates, and fully randomized within plates. Each plate contained: i) 87 samples from set 1 or set 2; ii) 1 empty well; iii) 2 Olink control samples used for quality control; iv) 3 Olink negative control samples used to compute the baseline assay level of each plate; and v) 3 Olink plate control samples used for normalization of protein expression. All Olink samples were in column 12 of each plate while the remaining 87 samples + 1 empty well were randomized across columns 1-11 and rows A to H. Plates were shipped into 8 batches (0-7) that consisted of different numbers of samples (Figure 1). Batches 0-6 contained exclusively samples from set 1, while batch 7 contained samples from both sets. A subset of 25 plates from batch 7 contained exclusively samples from set 1, while 94 of them contained samples from both sets.

**Figure S1: Schematic representation of the sample batch design for UKB.**

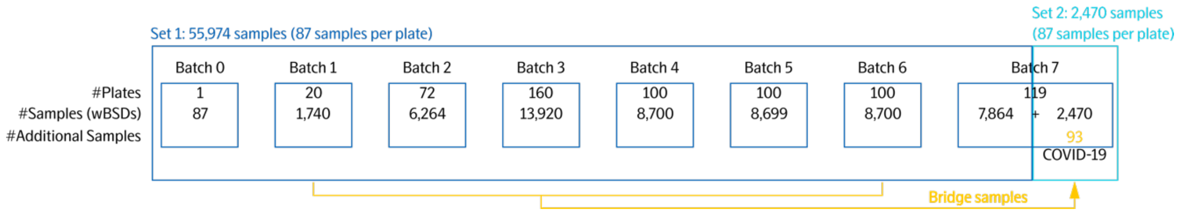

Calculation of Normalized Protein eXpression (NPX) values was performed stepwise as described below. Initially, we calculated the $log_2$ ratio of counts of each assay of each sample to the counts of the extension control, and the assay-specific median value of the plate controls was subtracted. This provided us with plate normalized NPX values for both sets. For samples belonging to set 1, we subtracted the assay-specific plate median NPX value, and we added the batch-specific median NPX value of each assay to account for effects within each batch. At this stage data was normalized within each batch. Next, we computed adjustment factors from the difference of the assay specific median NPX value of each batch to the reference batch (batch 1). Finally, adjustment factors from above were added to the NPX values of each batch of set 1. In summary, set 1 was normalized using a two-step approach of within-batch and across-batches intensity normalization. Samples of set 2 were normalized using reference (bridge) samples that were shared between the two sets. All plates that contained at least one sample of set 2 were assigned a randomly selected sample from the set 1. 93 samples with missing frequency <10% and representative of the NPX dynamic range were selected from batches 1-6 of set 1. These samples were assigned to the empty well of the 93 plates from batch 7 of set 2. In this case, adjustment factors were computed from the assay-specific median of the pair-wise differences between set 1 and set 2. Adjustment factors were added to the NPX values of set 2. Intensity normalized NPX values for set 1 and bridge normalized NPX values for set 2 consisted of the final set of NPX values that was used for the downstream analysis.

#### Data pre-processing and quality checking

The raw UKB-Olink data contained 58,699 samples and 54,309 individuals; taking out control and unprocessed samples and participants who have withdrawn from the study left 58,362 samples and 54,306 individuals. Further excluding data with QC failures (i.e., missing in NPX) led to 58,360 samples, and 54,304 individuals.

Outliers were identified by two approaches applied to each panel of proteins: (1) principal component analysis (PCA), and (2) examining the median and IQR of NPX across proteins by sample. We removed data points with (1) a standardized PC1 (the component that captures the most variation) or PC2 (second largest component) value more than 5 standard deviations from the mean (which is zero in standardized PCA), or (2) a median NPX greater than 5 standard deviations from the mean median, or an IQR of NPX greater than 5 standard deviations from the mean IQR.

After excluding outliers, we removed data points with a QC or assay warning, resulting 58,240 samples and 54,189 individuals in the data. We did not remove data based on LOD (lower limit of detection). The attrition of data, from excluding control sample to removing QC/assay warnings, is summarized in **Table S1**:

**Table S1. Summary of data attrition during QC.**

| | No. of samples | No. of individuals |
|---|---|---|
| **Raw data** | 58,699 | 54,309 |
| **Post-removal of control, unprocessed, withdrawn samples** | 58,362 | 54,306 |
| **Remove QC failures (i.e., missing in NPX)** | 58,360 | 54,304 |
| **Remove outliers** | 58,249 | 54,197 |
| **Remove QC or assay warnings** | 58,240 | 54,189 |

#### NPX data quality control

##### *Protein coefficients of variation (CVs)*

Two sets of duplicate samples were provided both by Olink Proteomics (‘Olink controls’) and UK Biobank (‘Blind Spike Duplications’ / BSDs). We measured the intra-person variability of protein $i$ of individual $j$ on plate $k$ for duplicate samples by calculating the coefficient of variation (CV) of NPX, as recommended by Olink:

$$CV_{ijk} = 100 * sqrt(\exp((log2 * SD(NPX))^2) - 1)$$

Then, to summarize the CV for each protein, we took the median of the CVs across individuals and plates by protein, plotting the median CVs for each protein:

$$median\_CV_i = median\left(CV_{ijk}\right) \text{ for each } j \text{ and } k$$

Intra-person CVs of each Olink protein are illustrated in **Figure S2**. The left panel of **Figure S2** shows that the CVs of the proteins range between 2.4% to 25%.

**Figure S2. Histograms of CVs (coefficients of variation) of proteins on the Olink platform**

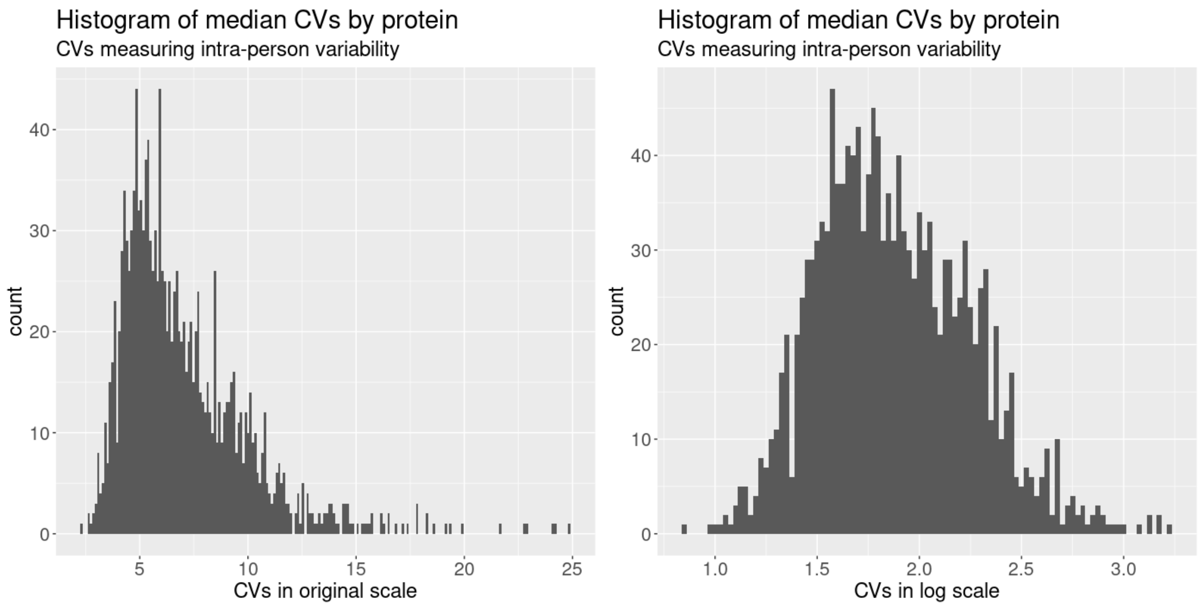

##### *Detecting batch effects*

To detect potential batch effects, we fit a simple random-effects model on each protein and calculated the percentage of variability attributable to each batch. A high percentage suggested greater variability in NPX across batch. The model used was:

$$NPX = b_0 + u_j + e_{jk} \text{ , where}$$

- $b_0$ = global mean NPX
- $u_j$ = batch random effects
- $e_{jk}$ = error terms
- Percentage of variability attributable to batch = $\frac{VAR(u_j)}{VAR(u_j+e_{jk})}$

The percentage of variability attributable to batch, by protein, is illustrated in **Figure S3.** Overall, the proportion of variability attributable to batch was low. No notable evidence of batch effects was observed.

**Figure S3. Percentage of variability attributable to batch by protein**

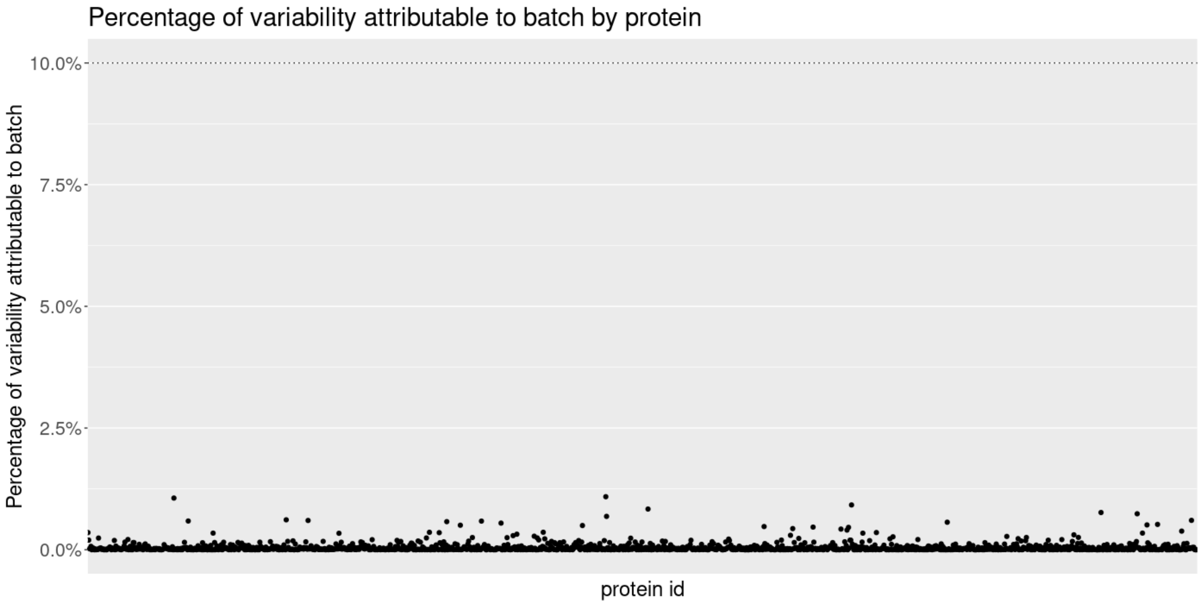

***Detecting plate effects***

We calculated two sets of CVs for each plate, plotting one series of CVs against the other, and identified whether any plate had high CVs on either axis of the plots. The two sets of CVs were defined as follows:

*Approach 1 - CVs measuring inter-replicate variability:*

1. Restrict the analysis to the Olink control samples, and calculate the CV by OlinkID and plate
2. Restrict the analysis to the BSD samples (same patient with two samples on the same plate), and calculate the CV by OlinkID and plate
3. Combine data from 1 and 2, and calculate the median of 1 and 2 by OlinkID and plate
4. Calculate the median across OlinkIDs by plate

*Approach 2 - CVs measuring inter-patient variability, including all patient samples but not Olink control samples):*

1. Calculate the CV by OlinkID and plate
2. Take the median CV across OlinkIDs by plate

We plotted the median CVs from approach 2 against approach 1. If any plate exhibited high CVs on either axis, we examined possible contributions towards the higher CV by:

1. Plotting the histograms of plate median CV, confirming that the plate(s) had a median CV that was not in range with others
2. Within the plate(s) with high CV, plotting the distribution of NPX by sample, to check whether the high median CV was driven by specific samples.

Inter-person versus inter-replicate variability for all plates is illustrated in **Figure S4**. Of all 600+ plates, only one showed evidence of abnormal outlying CVs – Plate 089000000130. The distribution of normalized protein expression levels by sample for this plate are illustrated in **Figure S5** and shows inter-replicate CV of plate 0890000000130 may be driven by a particular control sample (bottom-right panel Figure S4). Once that control sample is excluded, the inter-replicate CV of plate 0890000000130 is in range with others (**Figure S6**), and we conclude that no evidence of plate effect is observed.

**Figure S4**. **Plotting CV measuring intra-person vs inter-replicate variability for all plates.**

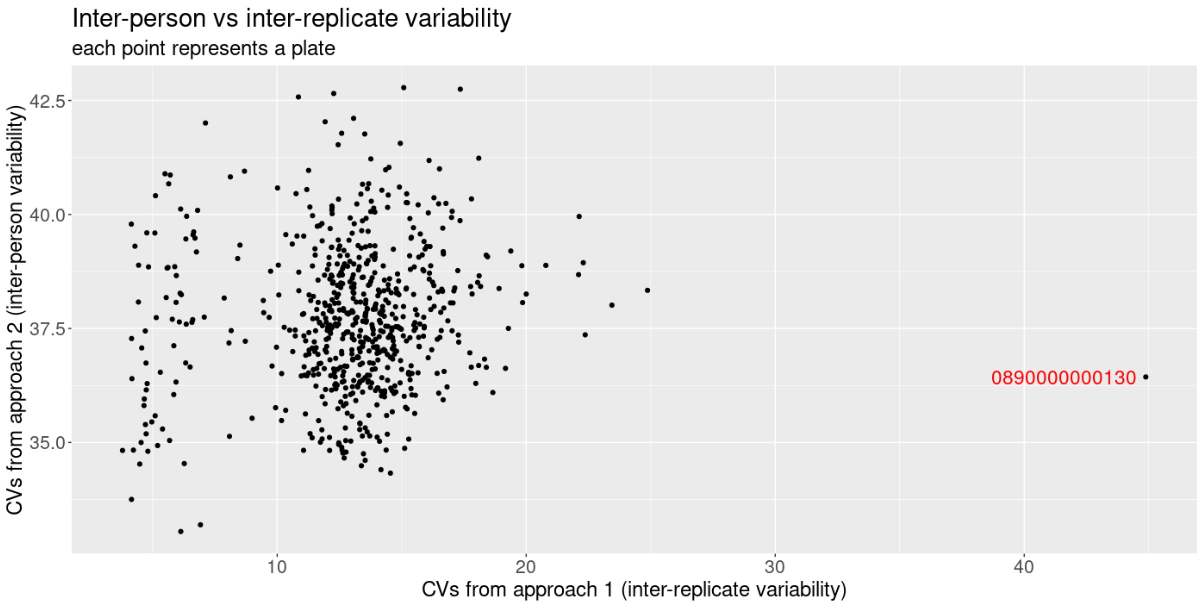

**Figure S5. Distribution of NPX by Sample for Plate 0890000000130**

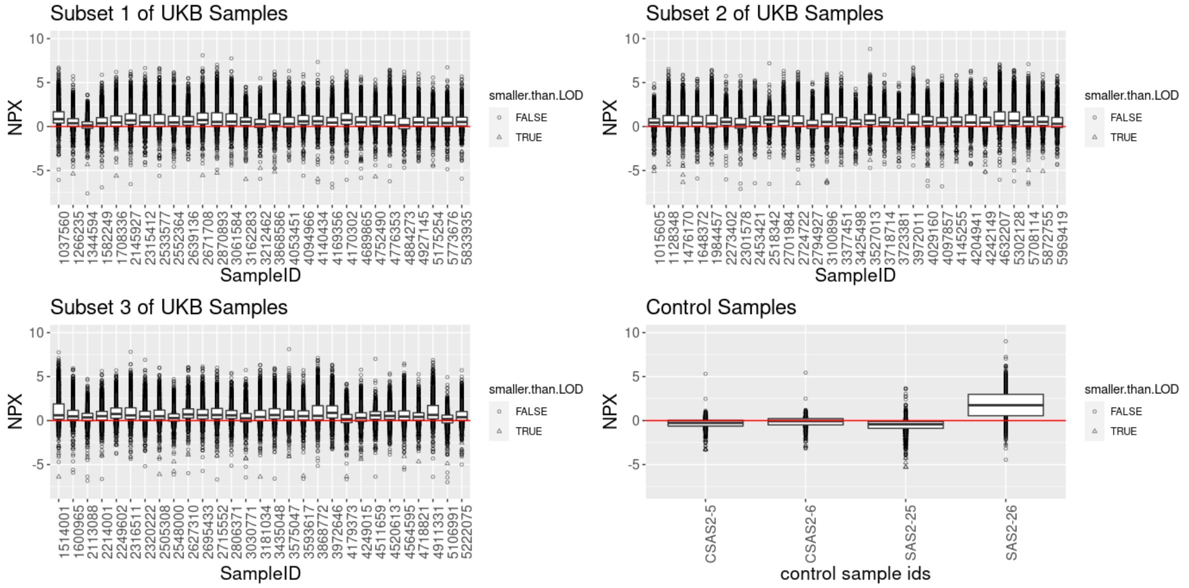

**Figure S6**. **Scatterplot of CV measuring inter-person vs inter-replicate variability after removing control sample SAS-26 from Plate 0890000000130**

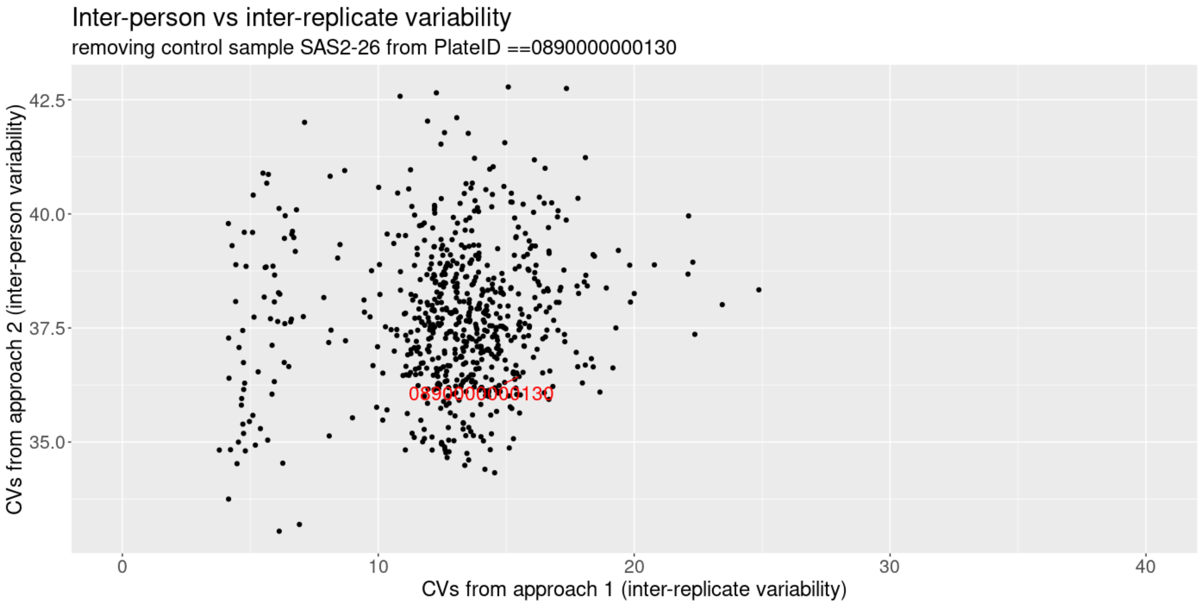

**Proteins below limit of detection (LOD)**

The proportion of samples with measurements below LOD for proteins across the panel are summarised in **Figure S7.** The vast majority of the protein analytes (1,328 out of 1,472) have the proportion of samples below LOD lower than 30%. 1,250 proteins have the proportion ranged between 0-10%; 44 and 34 proteins ranged between 10-20% and 20-30%, respectively. Among those with the proportion greater than 30%, 24 proteins are in the cardiometabolic panel; 39, 49 and 32 proteins in are the inflammation, neurology and oncology panel, respectively.

**Figure S7. Proportions of Samples with NPX measurements below LOD by protein panel**

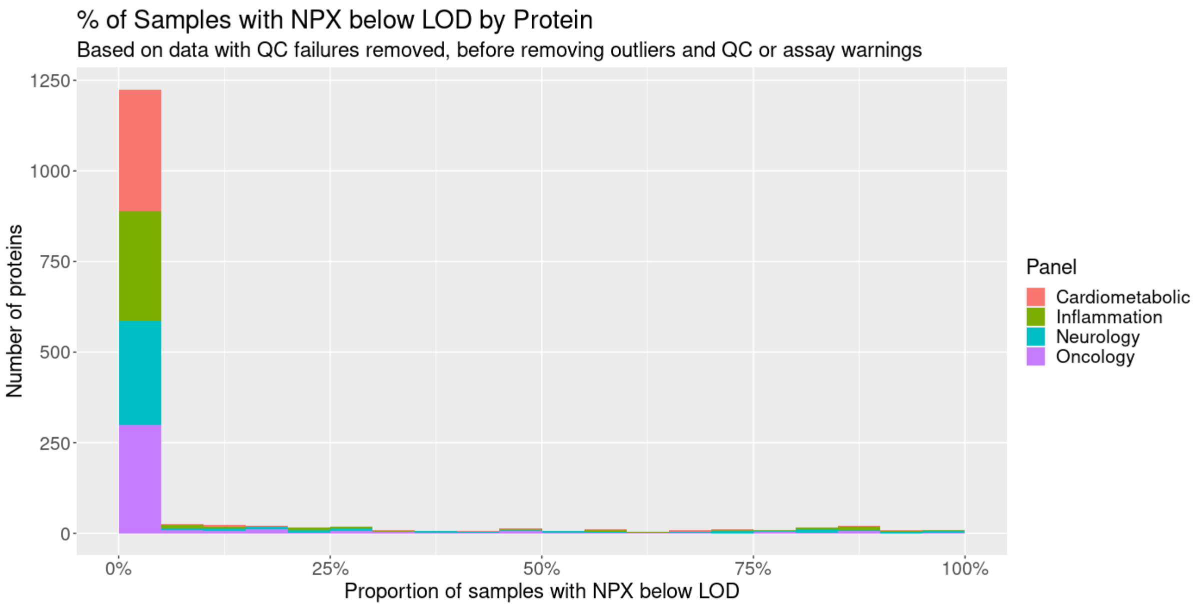

#### List of previous pQTL studies

To evaluate whether the pQTLs in the discovery set were novel, we used a list of published GWAS with proteomics (http://www.metabolomix.com/a-table-of-all-published-gwas-with-proteomics/) and the GWAS catalog to identify previously published pQTL studies. Twenty-six studies were included (**Table S2**). Using a *p*-value threshold of $3.4\text{x}10^{-11}$, we identified the sentinel variants and associated protein in the previously published studies and queried those against our discovery pQTLs. If a previously associated sentinel variant-protein pair fell within a 1 Mb window of the discovery set pQTL sentinel variant for the same protein and had an $r^2>0.8$ with any significant SNPs in the region, it was considered a replication.

**Table S2. List of previously published pQTL studies evaluated**

| Author | Publication date | PMID | URL | Proteomics technology |
|---|---|---|---|---|
| Melzer D | 2008-05-09 | 18464913 | https://pubmed.ncbi.nlm.nih.gov/18464913/ | immunoassay |
| Johansson A | 2013-03-19 | 23487758 | https://pubmed.ncbi.nlm.nih.gov/23487758/ | mass spectrometry |
| Kim S | 2013-07-23 | 23894628 | https://pubmed.ncbi.nlm.nih.gov/23894628/ | immunoassay |
| Enroth S | 2014-08-22 | 25147954 | https://pubmed.ncbi.nlm.nih.gov/25147954/ | PEA/OLINK |
| Kauwe JS | 2014-10-23 | 25340798 | https://pubmed.ncbi.nlm.nih.gov/25340798/ | immunoassay |
| Sun W | 2016-08-17 | 27532455 | https://pubmed.ncbi.nlm.nih.gov/27532455/ | immunoassay |
| Ahola-Olli AV | 2016-12-13 | 27989323 | https://pubmed.ncbi.nlm.nih.gov/27989323/ | immunoassay |
| Sasayama D | 2016-11-26 | 28031287 | https://pubmed.ncbi.nlm.nih.gov/28031287/ | Aptamer/Somascan |
| Suhre K | 2017-02-27 | 28240269 | https://pubmed.ncbi.nlm.nih.gov/28240269/ | Aptamer/Somascan |
| Deming Y | 2017-02-28 | 28247064 | https://pubmed.ncbi.nlm.nih.gov/28247064/ | immunoassay |
| Folkersen L | 2017-04-03 | 28369058 | https://pubmed.ncbi.nlm.nih.gov/28369058/ | PEA/OLINK |
| Carayol J | 2017-12-12 | 29234017 | https://pubmed.ncbi.nlm.nih.gov/29234017/ | Aptamer/Somascan |
| Sun BB | 2018-06-06 | 29875488 | https://pubmed.ncbi.nlm.nih.gov/29875488/ | Aptamer/Somascan |
| Emilsson V | 2018-08-02 | 30072576 | https://pubmed.ncbi.nlm.nih.gov/30072576/ | Aptamer/Somascan |
| Yao C | 2018-08-15 | 30111768 | https://pubmed.ncbi.nlm.nih.gov/30111768/ | immunoassay |
| Hillary RF | 2019-07-18 | 31320639 | https://pubmed.ncbi.nlm.nih.gov/31320639/ | PEA/OLINK |
| Ruffieux H | 2020-06-03 | 32492067 | https://pubmed.ncbi.nlm.nih.gov/32492067/ | Aptamer/Somascan |
| Zhong W | 2020-06-23 | 32576278 | https://pubmed.ncbi.nlm.nih.gov/32576278/ | PEA/OLINK |
| Bretherick A | 2020-07-06 | 32628676 | https://pubmed.ncbi.nlm.nih.gov/32628676/ | PEA/OLINK |
| Hillary RF | 2020-07-08 | 32641083 | https://pubmed.ncbi.nlm.nih.gov/32641083/ | PEA/OLINK |
| Folkersen L | 2020-10-16 | 33067605 | https://pubmed.ncbi.nlm.nih.gov/33067605/ | PEA/OLINK |
| Pietzner M | 2021-11-12 | 34648354 | https://pubmed.ncbi.nlm.nih.gov/34648354/ | Aptamer/Somascan |
| Ferkingstad E | 2021-02-21 | 34857953 | https://pubmed.ncbi.nlm.nih.gov/34857953/ | Aptamer/Somascan |
| Katz D | 2021-11-24 | 34814699 | https://pubmed.ncbi.nlm.nih.gov/34814699/ | Aptamer/Somascan |
| Png G | 2021-12-02 | 34857772 | https://pubmed.ncbi.nlm.nih.gov/34857772/ | PEA/OLINK |
| Gudjonsson A | 2022-01-25 | 35078996 | https://pubmed.ncbi.nlm.nih.gov/35078996/ | Aptamer/Somascan |

#### Sensitivity analyses of pQTLs

We also explored, *a priori,* the impact of blood cell composition, BMI, seasonal and fasting time before blood collection on pQTL effects (**Supplementary Table 17, Extended Data Figure 9**).

##### *Effects of blood cell counts*

Most primary associations from the discovery analysis (87.0% [8,912/10,248], including 99.8% [1,161/1,163] of *cis* and 85.3% [7,751/9,085] of *trans* associations) remained significant ($p<3.4\text{x}10^{-11}$) in the blood-cell sensitivity analysis, which adjusted for blood-cell composition (Methods). 79 *trans* associations (0.87% [79/9,085]) fell above nominal significance ($p>0.05$) with the addition of blood-cell covariates. Of these 79 *trans* associations, 75 were with a sentinel variant rs1354034 in *ARHGEF3*. In total, the *ARHGEF3* variant rs1354034 significantly associated with 239 proteins in the discovery analysis, and 60 proteins in the blood-cell sensitivity analysis. The *ARHGEF3* locus is established to be highly pleiotropic and known to associate with platelet counts[5,6]. A previous plasma pQTL study suggested that the observed pleiotropy at *ARHGEF3* may be driven by genetically determined increases in platelet counts and related sequelae that cause proteins to be secreted into plasma during sample handling and preparation[5]. We further tested this hypothesis through a formal mediation analysis of 179 variant – protein associations for platelet counts at the *ARHGEF3* locus using individual participant data (Methods). After correction for multiple testing (Bonferroni correction for 179 variant – protein associations and 9 blood cell phenotypes; $p=3.1\text{x}10^{-5}$), 82% of the associations were found to be fully mediated by platelet counts, in line with the observed decrease in pleiotropy at *ARHGEF3* variant rs1354034 after adjusting for blood-cell composition.

##### *Effects of BMI*

In the BMI-adjusted analysis, all but 2.0% [210/10,248; all *trans* associations] of the primary associations from the discovery analysis remained significant ($p<3.4\text{x}10^{-11}$). Of these 210 *trans* associations, only one association (leptin levels - rs56094641) fell below a significance threshold of $p<0.05$. This association with leptin levels was driven by an intronic variant (rs56094641) in *FTO*, an established obesity associated locus, suggesting that leptin association with this variant is mediated by obesity[7].

***Effects of season and amount of time fasted at blood collection***

The majority (99.5% [10,198/10,248]) of sentinel pQTLs identified by the discovery analysis remained genome-wide significant ($p<3.4\times10^{-11}$) after adjustment for season and participant-reported fasting time at blood collection. *P*-values for the 50 associations (all *trans* pQTLs) that were no longer genome-wide significant after accounting for multiple testing ranged from $3.43\times10^{-11}$ to $2.04\times10^{-10}$, suggesting minimal impact of season and/or fasting time on variant associations with protein levels.

#### Inflammasome – *NLRP12* locus

Previous proteo-genomics studies using the SomaScan aptamer platform reported inconsistent *trans* pQTL results at the *NLRP12* locus. The INTERVAL[8] study identified this as a pleotropic hot spot, whereas AGES and a cross-platform study did not[9,10], leading to speculation that pQTL associations at this locus may be platform dependent and/or the result of inter-study differences in sample handling.

Additional investigation of the independent UKB-PPP cohort suggest the associations observed here can reflect underlying biological genotype-dependent stress response, possibly as a result of sample perturbations, rather than technical artefacts. The INTERVAL study identified more than 300 *trans* pQTLs within a 5Kb window of the *NLRP12* locus. Using similar filtering criteria, we identified 26 *trans* pQTLs within the same window. While head-to-head comparisons between these studies are difficult due to greater statistical power of PPP and broader protein screening of INTERVAL, these results suggest the UKB-PPP findings are more consistent with the typical pleotropic loci identified in other studies.

In contrast to other loci such as *ARHGEF3* (see Effects of blood cell counts section), *trans* pQTL results at the *NLRP12* locus were robust to cell blood count sensitivity analyses. This supports the hypotheses that these signals result from inflammasome driven biological effects, rather than artificial increases in protein concentrations caused by sample handling related cell lysis.

### UKB-PPP consortium banner contributors

| Institute | Name | Address |
|---|---|---|
| Alnylam Human Genetics | Aimee M Deaton | Alnylam Human Genetics, Discovery & Translational Research, Alnylam Pharmaceuticals, Cambridge, MA, US |
| Alnylam Human Genetics | Rachel Hoffing | Alnylam Human Genetics, Discovery & Translational Research, Alnylam Pharmaceuticals, Cambridge, MA, US |
| Alnylam Human Genetics | Aaron M Holleman | Alnylam Human Genetics, Discovery & Translational Research, Alnylam Pharmaceuticals, Cambridge, MA, US |
| Alnylam Human Genetics | Lynn Krohn | Alnylam Human Genetics, Discovery & Translational Research, Alnylam Pharmaceuticals, Cambridge, MA, US |
| Alnylam Human Genetics | Philip LoGerfo | Alnylam Human Genetics, Discovery & Translational Research, Alnylam Pharmaceuticals, Cambridge, MA, US |
| Alnylam Human Genetics | Paul Nioi | Alnylam Human Genetics, Discovery & Translational Research, Alnylam Pharmaceuticals, Cambridge, MA, US |
| Alnylam Human Genetics | Mollie E Plekan | Alnylam Human Genetics, Discovery & Translational Research, Alnylam Pharmaceuticals, Cambridge, MA, US |
| Alnylam Human Genetics | Lucas D Ward | Alnylam Human Genetics, Discovery & Translational Research, Alnylam Pharmaceuticals, Cambridge, MA, US |
| Alnylam Human Genetics | Carissa Willis | Alnylam Human Genetics, Discovery & Translational Research, Alnylam Pharmaceuticals, Cambridge, MA, US |
| AstraZeneca Genomics Initiative | Bastian Angermann | Translational Science and Experimental Medicine, Research and Early Development, Respiratory and Immunology, BioPharmaceuticals R&D, AstraZeneca, Gothenburg, Sweden |
| AstraZeneca Genomics Initiative | Oliver Burren | Centre for Genomics Research, Discovery Sciences, BioPharmaceuticals R&D, AstraZeneca, Cambridge, UK |
| AstraZeneca Genomics Initiative | Keren Carss | Centre for Genomics Research, Discovery Sciences, BioPharmaceuticals R&D, AstraZeneca, Cambridge, UK |
| AstraZeneca Genomics Initiative | Ryan Dhindsa | Centre for Genomics Research, Discovery Sciences, BioPharmaceuticals R&D, AstraZeneca, Cambridge, USA |
| AstraZeneca Genomics Initiative | Ian Henry | Translational Science & Experimental Medicine, Research and Early Development, Cardiovascular, Renal and Metabolism, BioPharmaceuticals R&D, AstraZeneca, Cambridge, UK |
| AstraZeneca Genomics Initiative | Ventzi Hristova | Centre for Genomics Research, Discovery Sciences, BioPharmaceuticals R&D, AstraZeneca, Gaithersburg, USA |
| AstraZeneca Genomics Initiative | Daniel Muthas | Translational Science and Experimental Medicine, Research and Early Development, Respiratory and Immunology, BioPharmaceuticals R&D, AstraZeneca, Gothenburg, Sweden |
| AstraZeneca Genomics Initiative | Erin Oerton | Centre for Genomics Research, Discovery Sciences, BioPharmaceuticals R&D, AstraZeneca, Cambridge, UK |
| AstraZeneca Genomics Initiative | Menelas N Pangalos | BioPharmaceuticals R&D, AstraZeneca, Cambridge, UK |
| AstraZeneca Genomics Initiative | Dirk Paul | Centre for Genomics Research, Discovery Sciences, BioPharmaceuticals R&D, AstraZeneca, Cambridge, UK |
| AstraZeneca Genomics Initiative | Slave Petrovski | Centre for Genomics Research, Discovery Sciences, BioPharmaceuticals R&D, AstraZeneca, Cambridge, UK |
| AstraZeneca Genomics Initiative | Adam Platt | Translational Science and Experimental Medicine, Research and Early Development, Respiratory and Immunology, BioPharmaceuticals R&D, AstraZeneca, Cambridge, UK |
| AstraZeneca Genomics Initiative | Bram Prins | Centre for Genomics Research, Discovery Sciences, BioPharmaceuticals R&D, AstraZeneca, Cambridge, UK |
| AstraZeneca Genomics Initiative | Ben Sidders | Bioinformatics and Data Science, Research and Early Development, Oncology R&D, AstraZeneca, Cambridge, UK |
| AstraZeneca Genomics Initiative | Katherine R Smith | Centre for Genomics Research, Discovery Sciences, BioPharmaceuticals R&D, AstraZeneca, Cambridge, UK |
| AstraZeneca Genomics Initiative | Quanli Wang | Centre for Genomics Research, Discovery Sciences, BioPharmaceuticals R&D, AstraZeneca, Cambridge, USA |
| AstraZeneca Genomics Initiative | Sebastian Wasilewski | Centre for Genomics Research, Discovery Sciences, BioPharmaceuticals R&D, AstraZeneca, Cambridge, UK |
| AstraZeneca Genomics Initiative | Eleanor Wheeler | Centre for Genomics Research, Discovery Sciences, BioPharmaceuticals R&D, AstraZeneca, Cambridge, UK |
| Biogen Biobank Team | Denis Baird | Research & Development, Biogen Inc., Cambridge, MA, US |
| Biogen Biobank Team | Paola Bronson | Research & Development, Biogen Inc., Cambridge, MA, US |

| | | |
|---|---|---|
| Biogen Biobank Team | Danai Chasioti | Research & Development, Biogen Inc., Cambridge, MA, US |
| Biogen Biobank Team | Chia-Yen Chen | Research & Development, Biogen Inc., Cambridge, MA, US |
| Biogen Biobank Team | Susan Eaton | Research & Development, Biogen Inc., Cambridge, MA, US |
| Biogen Biobank Team | Amanda Edwards | Research & Development, Biogen Inc., Cambridge, MA, US |
| Biogen Biobank Team | Kyle Ferber | Research & Development, Biogen Inc., Cambridge, MA, US |
| Biogen Biobank Team | Jake Gagnon | Research & Development, Biogen Inc., Cambridge, MA, US |
| Biogen Biobank Team | Feng Gao | Research & Development, Biogen Inc., Cambridge, MA, US |
| Biogen Biobank Team | Cynthia Gubbels | Research & Development, Biogen Inc., Cambridge, MA, US |
| Biogen Biobank Team | Yunfeng Huang | Research & Development, Biogen Inc., Cambridge, MA, US |
| Biogen Biobank Team | Megan Jensen | Research & Development, Biogen Inc., Cambridge, MA, US |
| Biogen Biobank Team | Sally John | Research & Development, Biogen Inc., Cambridge, MA, US |
| Biogen Biobank Team | Varant Kupelian | Research & Development, Biogen Inc., Cambridge, MA, US |
| Biogen Biobank Team | Kejie Li | Research & Development, Biogen Inc., Cambridge, MA, US |
| Biogen Biobank Team | Stephanie Loomis | Research & Development, Biogen Inc., Cambridge, MA, US |
| Biogen Biobank Team | Eric Marshall | Research & Development, Biogen Inc., Cambridge, MA, US |
| Biogen Biobank Team | Helen McLaughlin | Research & Development, Biogen Inc., Cambridge, MA, US |
| Biogen Biobank Team | Adele Mitchell | Research & Development, Biogen Inc., Cambridge, MA, US |
| Biogen Biobank Team | Mehool Patel | Research & Development, Biogen Inc., Cambridge, MA, US |
| Biogen Biobank Team | Heiko Runz | Research & Development, Biogen Inc., Cambridge, MA, US |
| Biogen Biobank Team | Benjamin B Sun | Research & Development, Biogen Inc., Cambridge, MA, US |
| Biogen Biobank Team | Ellen Tsai | Research & Development, Biogen Inc., Cambridge, MA, US |
| Biogen Biobank Team | Christopher D Whelan | Research & Development, Biogen Inc., Cambridge, MA, US |
| Bristol Myers Squibb | Ian Catlett | Research and Early Development, BMS, Princeton, NJ, US |
| Bristol Myers Squibb | Ashok Dongre | Research and Early Development, BMS, Cambridge, MA, US |
| Bristol Myers Squibb | Joseph Maranville | Research and Early Development, BMS, Cambridge, MA, US |
| Bristol Myers Squibb | Peter Schafer | Research and Early Development, BMS, Princeton, NJ, US |
| Bristol Myers Squibb | Tai Wang | Research and Early Development, BMS, Cambridge, MA, US |
| Genentech Human Genetics | Christian Benner | Genentech, South San Francisco, CA, US |
| Genentech Human Genetics | Jerome Irudayanathan | Genentech, South San Francisco, CA, US |
| Genentech Human Genetics | Anubha Mahajan | Genentech, South San Francisco, CA, US |
| Genentech Human Genetics | Mark I McCarthy | Genentech, South San Francisco, CA, US |
| Genentech Human Genetics | Rion K Pendergrass | Genentech, South San Francisco, CA, US |
| Genentech Human Genetics | Xiaoman Xie | Genentech, South San Francisco, CA, US |
| GlaxoSmithKline Genomic Sciences | Elena Arciero | Genomic Sciences, GlaxoSmithKline, Stevenage, UK, UK |
| GlaxoSmithKline Genomic Sciences | Joanna C. Betts | Genomic Sciences, GlaxoSmithKline, Stevenage, UK, UK |
| GlaxoSmithKline Genomic Sciences | Nicholas Bowker | Genomic Sciences, GlaxoSmithKline, Stevenage, UK, UK |
| GlaxoSmithKline Genomic Sciences | Audrey Y. Chu | Genomic Sciences, GlaxoSmithKline, Collegeville, PA, US |
| GlaxoSmithKline Genomic Sciences | Adrian Cortes | Genomic Sciences, GlaxoSmithKline, Stevenage, UK, UK |
| GlaxoSmithKline Genomic Sciences | Damien C. Croteau-Chonka | Genomic Sciences, GlaxoSmithKline, Collegeville, PA, US |
| GlaxoSmithKline Genomic Sciences | Emmanouil T. Dermitzakis | Genomic Sciences, GlaxoSmithKline, Collegeville, PA, US |
| GlaxoSmithKline Genomic Sciences | Margaret G. Ehm | Genomic Sciences, GlaxoSmithKline, Collegeville, PA, US |
| GlaxoSmithKline Genomic Sciences | Stephan Gade | Genomic Sciences, GlaxoSmithKline, Heidelberg, Germany |
| GlaxoSmithKline Genomic Sciences | Padhraig Gormley | Genomic Sciences, GlaxoSmithKline, Collegeville, PA, US |
| GlaxoSmithKline Genomic Sciences | Erik D. Ingelsson | Genomic Sciences, GlaxoSmithKline, Collegeville, PA, US |
| GlaxoSmithKline Genomic Sciences | Toby Johnson | Genomic Sciences, GlaxoSmithKline, Stevenage, UK, UK |
| GlaxoSmithKline Genomic Sciences | Jimmy Zhenli Liu | Genomic Sciences, GlaxoSmithKline, Collegeville, PA, US |

| | | |
|---|---|---|
| GlaxoSmithKline Genomic Sciences | Yancy Lo | Genomic Sciences, GlaxoSmithKline, Collegeville, PA, US |
| GlaxoSmithKline Genomic Sciences | Jatin Sandhuria | Genomic Sciences, GlaxoSmithKline, Stevenage, UK, UK |
| GlaxoSmithKline Genomic Sciences | Richard M. Turner | Genomic Sciences, GlaxoSmithKline, Stevenage, UK, UK |
| GlaxoSmithKline Genomic Sciences | Qin Wang | Genomic Sciences, GlaxoSmithKline, Stevenage, UK, UK |
| GlaxoSmithKline Genomic Sciences | Frederik Ziebell | Genomic Sciences, GlaxoSmithKline, Heidelberg, Germany |
| Pfizer Integrative Biology | Xinli Hu | Inflammation and Immunology Research Unit, Worldwide Research, Development and Medical, Pfizer, Cambridge, MA, US |
| Pfizer Integrative Biology | Craig L. Hyde | Non-Clinical Research Statistics, Early Clinical Development, Worldwide Research, Development and Medical, Pfizer, Groton, CT, US |
| Pfizer Integrative Biology | Hye In Kim | Internal Medicine Research Unit, Worldwide Research, Development and Medical, Pfizer, Cambridge, MA, US |
| Pfizer Integrative Biology | A. Katrina Loomis | External Science and Innovation Target Sciences, Worldwide Research, Development and Medical, Pfizer, Groton, CT, US |
| Pfizer Integrative Biology | Anders Malarstig | External Science and Innovation Target Sciences, Worldwide Research, Development and Medical, Pfizer, Stockholm, Sweden |
| Pfizer Integrative Biology | Zhan Ye | Non-Clinical Research Statistics, Early Clinical Development, Worldwide Research, Development and Medical, Pfizer, Cambridge, MA, US |
| Population Analytics of Janssen Data Sciences | Evan H Baugh | Population Analytics, Data Sciences, Janssen R&D, Spring House, PA, US |
| Population Analytics of Janssen Data Sciences | Mary Helen Black | Population Analytics, Data Sciences, Janssen R&D, Spring House, PA, US |
| Population Analytics of Janssen Data Sciences | Abolfazl Doostparast Torshizi | Population Analytics, Data Sciences, Janssen R&D, Spring House, PA, US |
| Population Analytics of Janssen Data Sciences | Shicheng Guo | Population Analytics, Data Sciences, Janssen R&D, Spring House, PA, US |
| Population Analytics of Janssen Data Sciences | Hussein A Hejase | Population Analytics, Data Sciences, Janssen R&D, Spring House, PA, US |
| Population Analytics of Janssen Data Sciences | Liping Hou | Population Analytics, Data Sciences, Janssen R&D, Spring House, PA, US |
| Population Analytics of Janssen Data Sciences | Shuwei Li | Population Analytics, Data Sciences, Janssen R&D, Spring House, PA, US |
| Population Analytics of Janssen Data Sciences | Alexander H Li | Population Analytics, Data Sciences, Janssen R&D, Spring House, PA, US |
| Population Analytics of Janssen Data Sciences | Brice AJ Sarver | Population Analytics, Data Sciences, Janssen R&D, Spring House, PA, US |
| Regeneron Genetics Center | Gonçalo Abecasis | Regeneron Genetics Center, Tarrytown, NY, US |
| Regeneron Genetics Center | Parsa Akbari | Regeneron Genetics Center, Tarrytown, NY, US |
| Regeneron Genetics Center | Anna Alkelai | Regeneron Genetics Center, Tarrytown, NY, US |
| Regeneron Genetics Center | Manuel Allen Revez Ferreira | Regeneron Genetics Center, Tarrytown, NY, US |
| Regeneron Genetics Center | Amelia Averitt | Regeneron Genetics Center, Tarrytown, NY, US |
| Regeneron Genetics Center | Ariane Ayer | Regeneron Genetics Center, Tarrytown, NY, US |
| Regeneron Genetics Center | Joshua Backman | Regeneron Genetics Center, Tarrytown, NY, US |
| Regeneron Genetics Center | Xiaodong Bai | Regeneron Genetics Center, Tarrytown, NY, US |
| Regeneron Genetics Center | Suganthi Balasubramanian | Regeneron Genetics Center, Tarrytown, NY, US |
| Regeneron Genetics Center | Nilanjana Banerjee | Regeneron Genetics Center, Tarrytown, NY, US |
| Regeneron Genetics Center | Suying Bao | Regeneron Genetics Center, Tarrytown, NY, US |
| Regeneron Genetics Center | Aris Baras | Regeneron Genetics Center, Tarrytown, NY, US |
| Regeneron Genetics Center | Christina Beechert | Regeneron Genetics Center, Tarrytown, NY, US |
| Regeneron Genetics Center | Boris Boutkov | Regeneron Genetics Center, Tarrytown, NY, US |
| Regeneron Genetics Center | Jonas Bovijn | Regeneron Genetics Center, Tarrytown, NY, US |
| Regeneron Genetics Center | Erin D Brian | Regeneron Genetics Center, Tarrytown, NY, US |
| Regeneron Genetics Center | Michael Cantor | Regeneron Genetics Center, Tarrytown, NY, US |
| Regeneron Genetics Center | Esteban Chen | Regeneron Genetics Center, Tarrytown, NY, US |

| | | |
|---|---|---|
| Regeneron Genetics Center | Siying Chen | Regeneron Genetics Center, Tarrytown, NY, US |
| Regeneron Genetics Center | Shek Man Chim | Regeneron Genetics Center, Tarrytown, NY, US |
| Regeneron Genetics Center | Shing Wan Choi | Regeneron Genetics Center, Tarrytown, NY, US |
| Regeneron Genetics Center | Janice Clauer | Regeneron Genetics Center, Tarrytown, NY, US |
| Regeneron Genetics Center | Thomas Coleman | Regeneron Genetics Center, Tarrytown, NY, US |
| Regeneron Genetics Center | Giovanni Coppola | Regeneron Genetics Center, Tarrytown, NY, US |
| Regeneron Genetics Center | Ruan Cox | Regeneron Genetics Center, Tarrytown, NY, US |
| Regeneron Genetics Center | Laura Cremona | Regeneron Genetics Center, Tarrytown, NY, US |
| Regeneron Genetics Center | Amy Damask | Regeneron Genetics Center, Tarrytown, NY, US |
| Regeneron Genetics Center | Tanima De | Regeneron Genetics Center, Tarrytown, NY, US |
| Regeneron Genetics Center | Giusy Della Gatta | Regeneron Genetics Center, Tarrytown, NY, US |
| Regeneron Genetics Center | Andrew Deubler | Regeneron Genetics Center, Tarrytown, NY, US |
| Regeneron Genetics Center | Alessandro Di Gioia | Regeneron Genetics Center, Tarrytown, NY, US |
| Regeneron Genetics Center | Lee Dobbyn | Regeneron Genetics Center, Tarrytown, NY, US |
| Regeneron Genetics Center | Valerio Donato | Regeneron Genetics Center, Tarrytown, NY, US |
| Regeneron Genetics Center | Peter Dornbos | Regeneron Genetics Center, Tarrytown, NY, US |
| Regeneron Genetics Center | Hang Du | Regeneron Genetics Center, Tarrytown, NY, US |
| Regeneron Genetics Center | Aris Economides | Regeneron Genetics Center, Tarrytown, NY, US |
| Regeneron Genetics Center | Gisu Eom | Regeneron Genetics Center, Tarrytown, NY, US |
| Regeneron Genetics Center | Alison Fenney | Regeneron Genetics Center, Tarrytown, NY, US |
| Regeneron Genetics Center | Daniel Fernandez | Regeneron Genetics Center, Tarrytown, NY, US |
| Regeneron Genetics Center | Adolfo Ferrando | Regeneron Genetics Center, Tarrytown, NY, US |
| Regeneron Genetics Center | Caitlin Forsythe | Regeneron Genetics Center, Tarrytown, NY, US |
| Regeneron Genetics Center | Jan Freudenberg | Regeneron Genetics Center, Tarrytown, NY, US |
| Regeneron Genetics Center | Sheila Gaynor | Regeneron Genetics Center, Tarrytown, NY, US |
| Regeneron Genetics Center | Sahar Gelfman | Regeneron Genetics Center, Tarrytown, NY, US |
| Regeneron Genetics Center | Benjamin Geraghty | Regeneron Genetics Center, Tarrytown, NY, US |
| Regeneron Genetics Center | Akropravo Ghosh | Regeneron Genetics Center, Tarrytown, NY, US |
| Regeneron Genetics Center | Christopher Gillies | Regeneron Genetics Center, Tarrytown, NY, US |
| Regeneron Genetics Center | Sujit Gokhale | Regeneron Genetics Center, Tarrytown, NY, US |
| Regeneron Genetics Center | Alexdander Gorovits | Regeneron Genetics Center, Tarrytown, NY, US |
| Regeneron Genetics Center | Sarah Graham | Regeneron Genetics Center, Tarrytown, NY, US |
| Regeneron Genetics Center | Zhenhua Gu | Regeneron Genetics Center, Tarrytown, NY, US |
| Regeneron Genetics Center | Kristy Guevara | Regeneron Genetics Center, Tarrytown, NY, US |
| Regeneron Genetics Center | Lauren Gurski | Regeneron Genetics Center, Tarrytown, NY, US |
| Regeneron Genetics Center | Aysegul Guvenek | Regeneron Genetics Center, Tarrytown, NY, US |
| Regeneron Genetics Center | Mary Haas | Regeneron Genetics Center, Tarrytown, NY, US |
| Regeneron Genetics Center | Lukas Habegger | Regeneron Genetics Center, Tarrytown, NY, US |
| Regeneron Genetics Center | Jody Hankins | Regeneron Genetics Center, Tarrytown, NY, US |
| Regeneron Genetics Center | Samnuel Hart | Regeneron Genetics Center, Tarrytown, NY, US |
| Regeneron Genetics Center | Alicia Hawes | Regeneron Genetics Center, Tarrytown, NY, US |
| Regeneron Genetics Center | Joseph Herman | Regeneron Genetics Center, Tarrytown, NY, US |
| Regeneron Genetics Center | George Hindy | Regeneron Genetics Center, Tarrytown, NY, US |
| Regeneron Genetics Center | Kristen Howell | Regeneron Genetics Center, Tarrytown, NY, US |
| Regeneron Genetics Center | Momodow Jallow | Regeneron Genetics Center, Tarrytown, NY, US |
| Regeneron Genetics Center | Marcus B Jones | Regeneron Genetics Center, Tarrytown, NY, US |
| Regeneron Genetics Center | Eric Jorgenson | Regeneron Genetics Center, Tarrytown, NY, US |
| Regeneron Genetics Center | Tyler Joseph | Regeneron Genetics Center, Tarrytown, NY, US |
| Regeneron Genetics Center | Amit Joshi | Regeneron Genetics Center, Tarrytown, NY, US |
| Regeneron Genetics Center | Hyun Min Kang | Regeneron Genetics Center, Tarrytown, NY, US |
| Regeneron Genetics Center | Manav Kapoor | Regeneron Genetics Center, Tarrytown, NY, US |
| Regeneron Genetics Center | Katia Karalis | Regeneron Genetics Center, Tarrytown, NY, US |
| Regeneron Genetics Center | Michael Kessler | Regeneron Genetics Center, Tarrytown, NY, US |
| Regeneron Genetics Center | Lori Khrimian | Regeneron Genetics Center, Tarrytown, NY, US |
| Regeneron Genetics Center | Minhee Kim | Regeneron Genetics Center, Tarrytown, NY, US |
| Regeneron Genetics Center | Alexandros Kokkosis | Regeneron Genetics Center, Tarrytown, NY, US |
| Regeneron Genetics Center | Jack Kosmicki | Regeneron Genetics Center, Tarrytown, NY, US |
| Regeneron Genetics Center | Olga Krasheninina | Regeneron Genetics Center, Tarrytown, NY, US |
| Regeneron Genetics Center | Rouel Lanche | Regeneron Genetics Center, Tarrytown, NY, US |

| | | |
|---|---|---|
| Regeneron Genetics Center | Michael Lattari | Regeneron Genetics Center, Tarrytown, NY, US |
| Regeneron Genetics Center | Michelle G LeBlanc | Regeneron Genetics Center, Tarrytown, NY, US |
| Regeneron Genetics Center | Michelle Leddy | Regeneron Genetics Center, Tarrytown, NY, US |
| Regeneron Genetics Center | Dadong Li | Regeneron Genetics Center, Tarrytown, NY, US |
| Regeneron Genetics Center | Nan Lin | Regeneron Genetics Center, Tarrytown, NY, US |
| Regeneron Genetics Center | Daren Liu | Regeneron Genetics Center, Tarrytown, NY, US |
| Regeneron Genetics Center | Adam Locke | Regeneron Genetics Center, Tarrytown, NY, US |
| Regeneron Genetics Center | Alexander Lopez | Regeneron Genetics Center, Tarrytown, NY, US |
| Regeneron Genetics Center | Luca A Lotta | Regeneron Genetics Center, Tarrytown, NY, US |
| Regeneron Genetics Center | Vrushall Mahajan | Regeneron Genetics Center, Tarrytown, NY, US |
| Regeneron Genetics Center | Sameer Malhotra | Regeneron Genetics Center, Tarrytown, NY, US |
| Regeneron Genetics Center | Kia Manoochehri | Regeneron Genetics Center, Tarrytown, NY, US |
| Regeneron Genetics Center | Adam J Mansfield | Regeneron Genetics Center, Tarrytown, NY, US |
| Regeneron Genetics Center | Jonathan Marchini | Regeneron Genetics Center, Tarrytown, NY, US |
| Regeneron Genetics Center | Anthony Marcketta | Regeneron Genetics Center, Tarrytown, NY, US |
| Regeneron Genetics Center | Hector Martinez | Regeneron Genetics Center, Tarrytown, NY, US |
| Regeneron Genetics Center | Evan K Maxwell | Regeneron Genetics Center, Tarrytown, NY, US |
| Regeneron Genetics Center | Joelle Mbatchou | Regeneron Genetics Center, Tarrytown, NY, US |
| Regeneron Genetics Center | Jason Mighty | Regeneron Genetics Center, Tarrytown, NY, US |
| Regeneron Genetics Center | Lawrence Miloscio | Regeneron Genetics Center, Tarrytown, NY, US |
| Regeneron Genetics Center | Lyndon J Mitnaul | Regeneron Genetics Center, Tarrytown, NY, US |
| Regeneron Genetics Center | George Mitra | Regeneron Genetics Center, Tarrytown, NY, US |
| Regeneron Genetics Center | George Mitra | Regeneron Genetics Center, Tarrytown, NY, US |
| Regeneron Genetics Center | Arden Moscati | Regeneron Genetics Center, Tarrytown, NY, US |
| Regeneron Genetics Center | Mona Nafde | Regeneron Genetics Center, Tarrytown, NY, US |
| Regeneron Genetics Center | Priyanka Nakka | Regeneron Genetics Center, Tarrytown, NY, US |
| Regeneron Genetics Center | Jonas Nielsen | Regeneron Genetics Center, Tarrytown, NY, US |
| Regeneron Genetics Center | Nirupama Nishtala | Regeneron Genetics Center, Tarrytown, NY, US |
| Regeneron Genetics Center | Sheilyn Nunez | Regeneron Genetics Center, Tarrytown, NY, US |
| Regeneron Genetics Center | Sean O'Keeffe | Regeneron Genetics Center, Tarrytown, NY, US |
| Regeneron Genetics Center | John D Overton | Regeneron Genetics Center, Tarrytown, NY, US |
| Regeneron Genetics Center | Maria Padilla | Regeneron Genetics Center, Tarrytown, NY, US |
| Regeneron Genetics Center | Michelle Pagan | Regeneron Genetics Center, Tarrytown, NY, US |
| Regeneron Genetics Center | Anita Pandit | Regeneron Genetics Center, Tarrytown, NY, US |
| Regeneron Genetics Center | Razvan Panea | Regeneron Genetics Center, Tarrytown, NY, US |
| Regeneron Genetics Center | Charles Paulding | Regeneron Genetics Center, Tarrytown, NY, US |
| Regeneron Genetics Center | Elias Pavlopoulos | Regeneron Genetics Center, Tarrytown, NY, US |
| Regeneron Genetics Center | Ann Perez-Beals | Regeneron Genetics Center, Tarrytown, NY, US |
| Regeneron Genetics Center | Trikaldarshi Persaud | Regeneron Genetics Center, Tarrytown, NY, US |
| Regeneron Genetics Center | Tommy Polanco | Regeneron Genetics Center, Tarrytown, NY, US |
| Regeneron Genetics Center | Manasi Pradhan | Regeneron Genetics Center, Tarrytown, NY, US |
