## Extended Data Figures for "Genetic regulation of the human plasma proteome in 54,306 UK Biobank participants"

**Extended Data Figure 1. Summary of the Olink Explore proteomics assay Olink.** (a) Summary of the Olink proteomic assay workflow. (i) Assays are run in a 96-well format, each plate consists of 88 UKB samples and 8 external control samples in column 12: sample controls (yellow) are used to determine precision within and between plates, triplicate negative controls samples (red) set the limit of detection (LOD) and triplicate plate controls (green) are used to standardize protein levels within a plate. The Explore 1536 product consists of four 384-panels, Cardiometabolic (CAR), Inflammation (INF), Neurology (NEU) and Oncology (ONC), and each panel consists of 4 abundance blocks, with plasma sample run 1:1 or diluted 1:1, 1:10, 1:100, 1;1000. (ii) Extension and amplification step: only matched PEA probes bind to their respective target and via PCR (PCR1) generate dsDNA amplicons, containing assay information. (iii) Indexing: all amplicons for a given sample in a single panel are pooled and unique index primers added and are integrated in to the amplicon via PCR (PCR2). (iv) All amplicons for all samples within a panel are combined to generate four sequencing libraries, and the libraries purified, quality controlled before (v) detection and being sequenced on an Illumina Novaseq 6000 instrument generating ~150,000 data points per sample plate (b) Cell compartment distribution of measured proteins by protein panel.

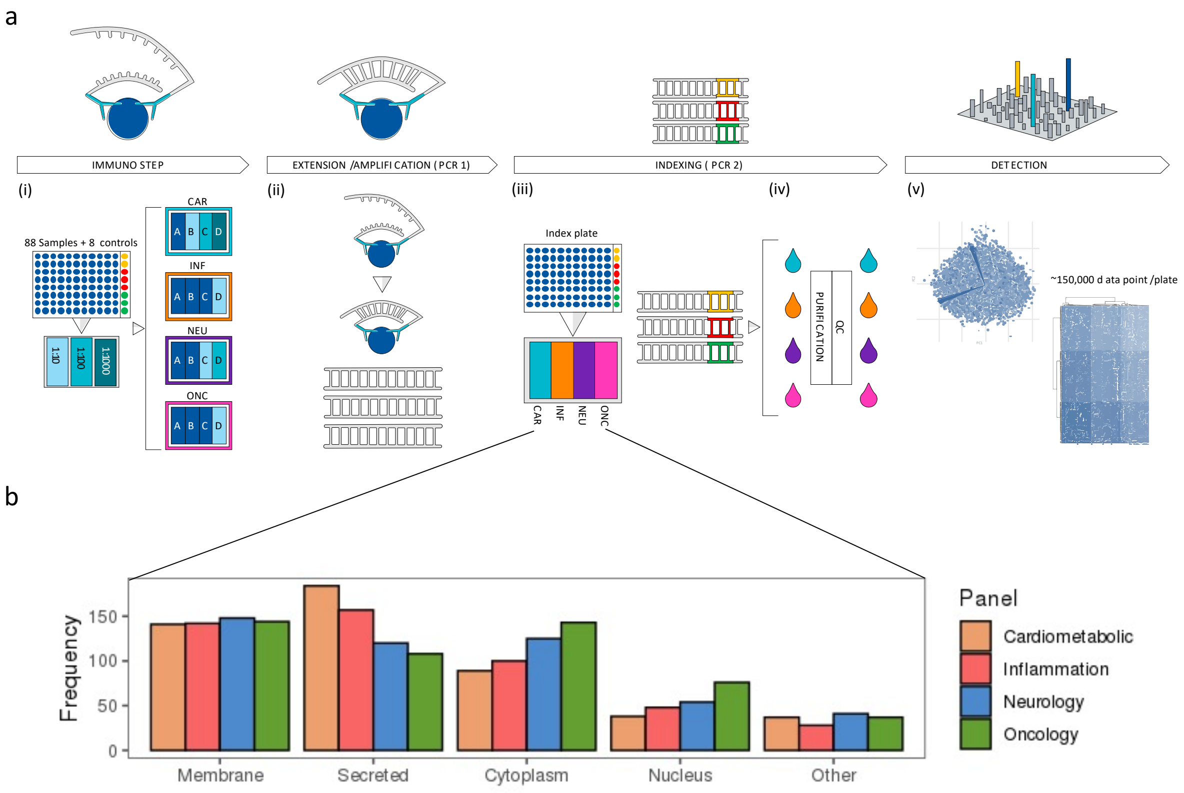

**Extended Data Figure 2.** (a) Phenotypic correlation (Pearson's *r*) between same protein targets (CXCL8, IL6, TNF) measured across all four protein panels. (b) Correlation (Pearson's *r*) of significant associations ($p<3.4\text{x}10^{-11}$) between the same protein targets.

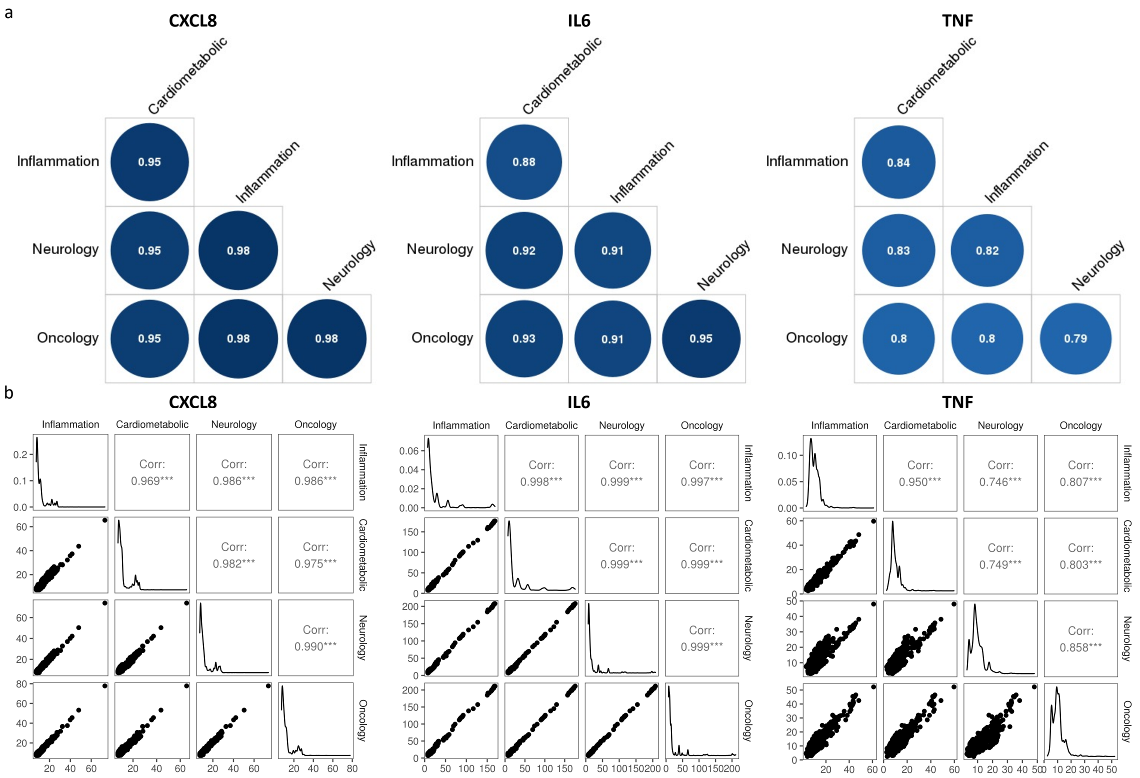

**Extended Data Figure 3.** (a) Volcano plot of associations with age, sex and BMI. Top 10 protein largest positive and negative associations labelled. (b) Comparison of effect sizes between UKB-PPP and INTERVAL (*Genomic atlas of the human plasma proteome, Nature 2018, Supplementary Table 2*) for protein associations with age, sex and BMI.
*r: Pearson's correlation coefficient.*

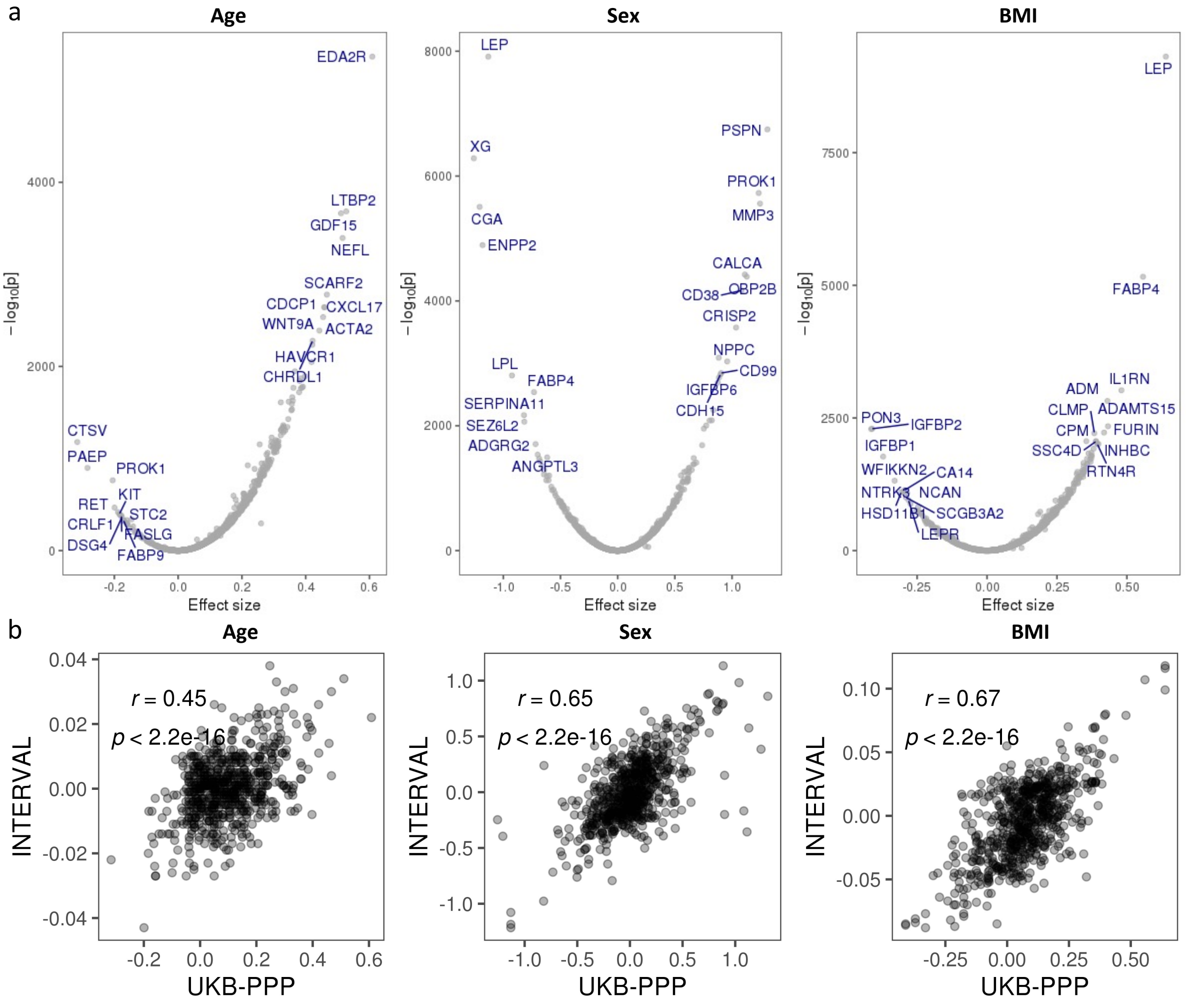

**Extended Data Figure 4.** (a) Comparison of number of pQTLs *vs* the proportion of samples with measurements below LOD for each protein. (b) Density plot of proportion of samples with measurements below LOD for proteins with no significant pQTLs ($p<3.4x10^{-11}$).
LOD: limit of detection. *ρ: Spearman's correlation coefficient.*

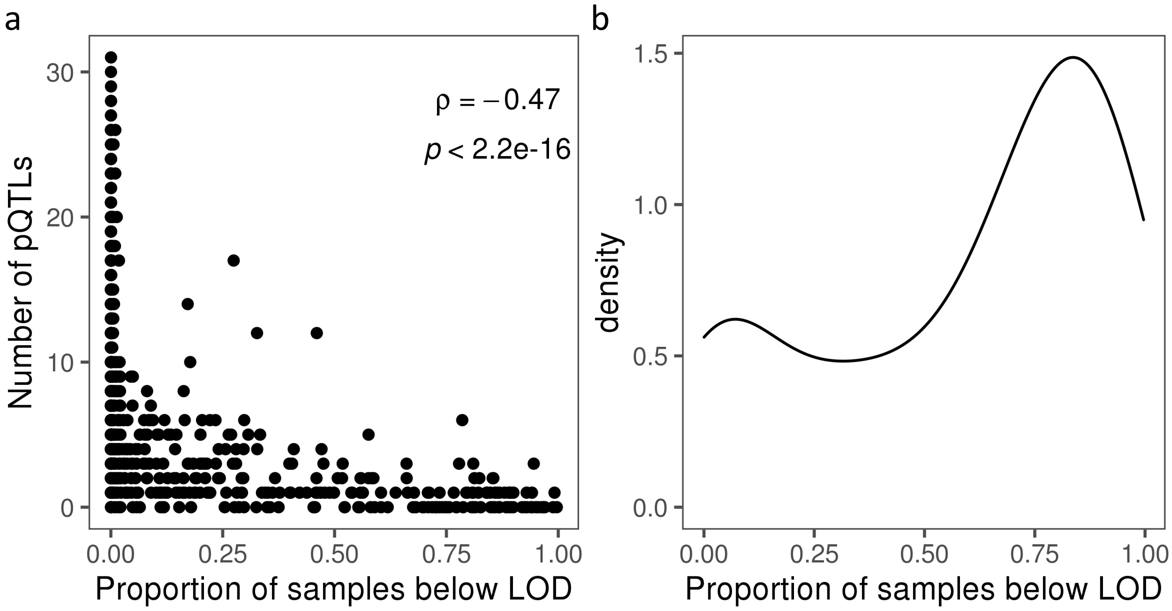

**Extended Data Figure 5.** (a) Annotation of primary pQTL variants. (b) SIFT and (c) PolyPhen annotations for primary pQTLs variants.

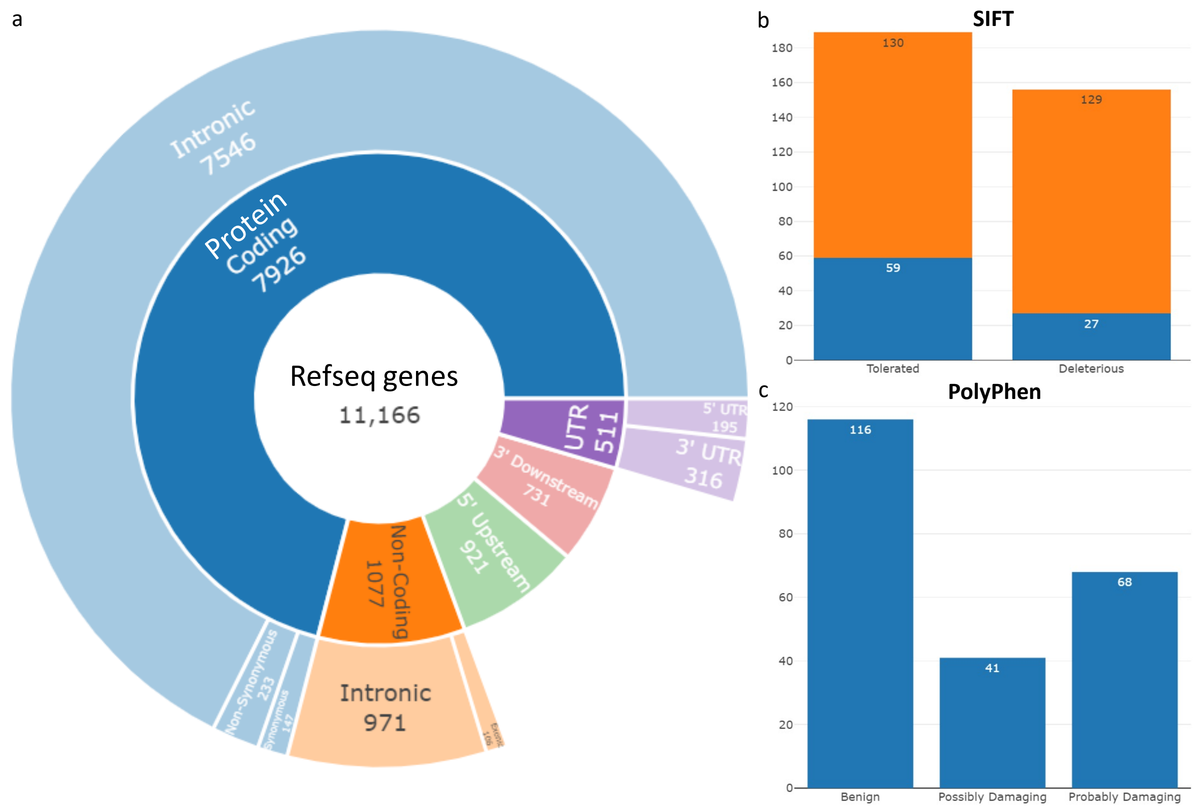

**Extended Data Figure 6.** (a) Comparison of effect sizes between discovery and replication cohorts. (b) Density plot of proportion of total heritability explained by primary *cis* and *trans* associations. (c) Scatterplot with overlaid regression line of the pQTL component (variance explained by sentinel primary pQTLs) *vs* the polygenic component (genome-wide SNP heritability excluding pQTL regions).
*ρ: Spearman's correlation coefficient.*

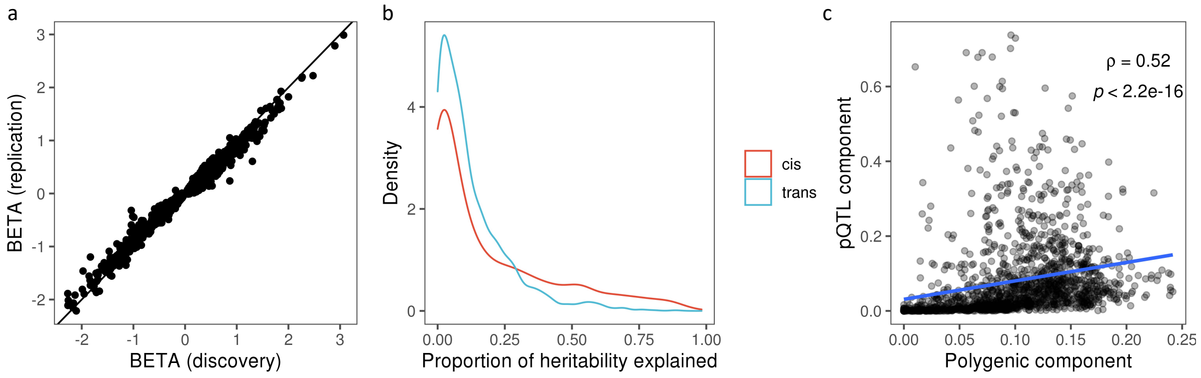

**Extended Data Figure 7.** Number of independent signals per region (a) and per protein (b).

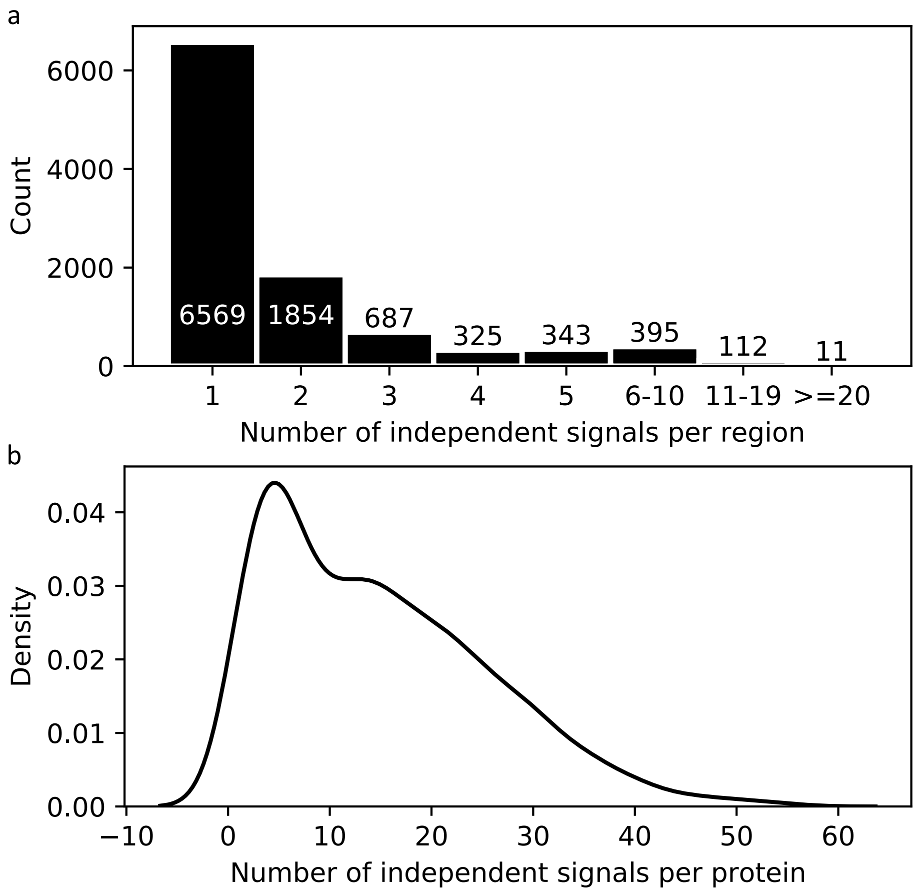

**Extended Data Figure 8.** (a) Number of proteins associated per genomic region at different sample sizes. (b) Number of proteins with at least one interaction partner loci (gene product at the *trans* locus that interacts with the protein tested) in at least one of the associated *trans* loci.

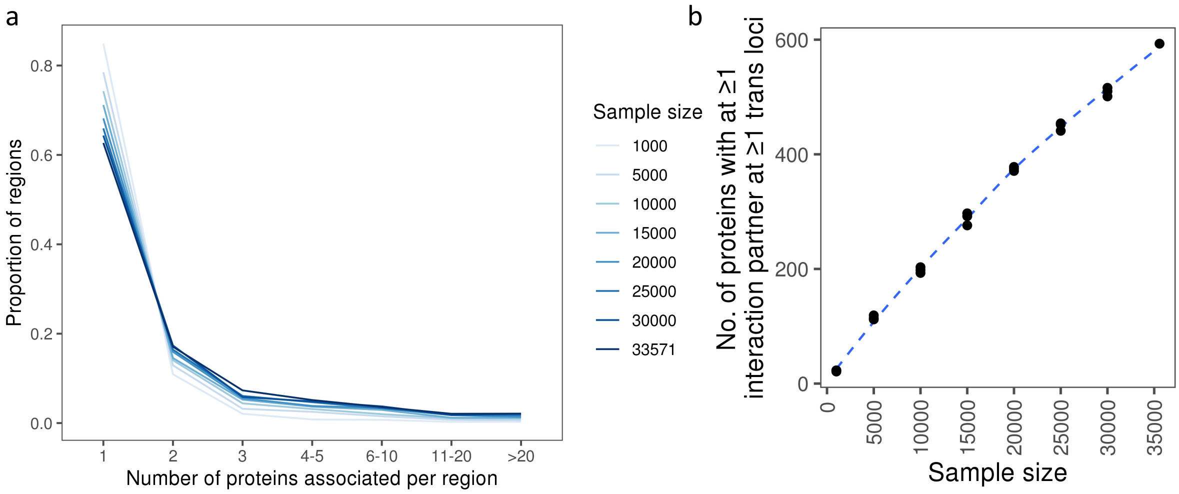

**Extended Data Figure 9. Comparison of effect size and p-value changes before and after sensitivity analyses.**

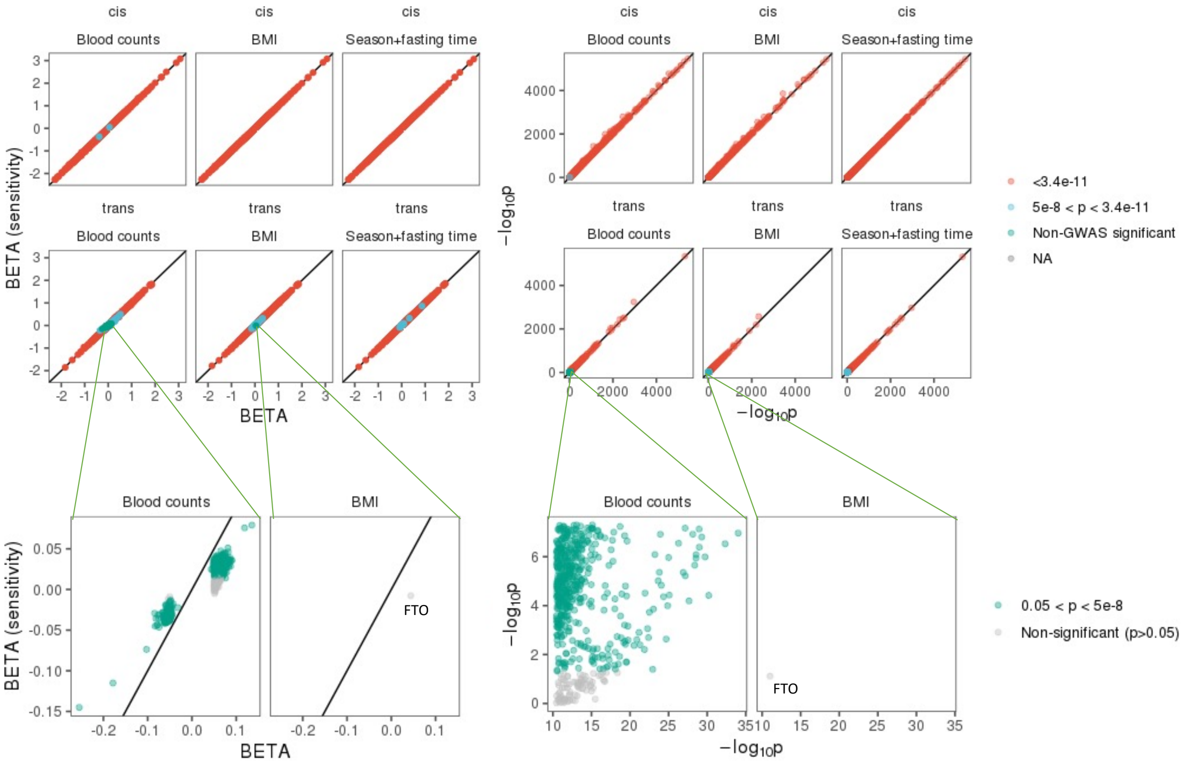

**Extended Data Figure 10. Summary of colocalization with eQTLs.** (a) Number of tissues with eQTL colocalizations across proteins (protein index on x axis). (b) Comparison of effect sizes between pQTL and eQTLs across multiple tissues. (c) Effects of the PARK7 pQTL on *PARK7* gene expression for colocalized tissue eQTLs.

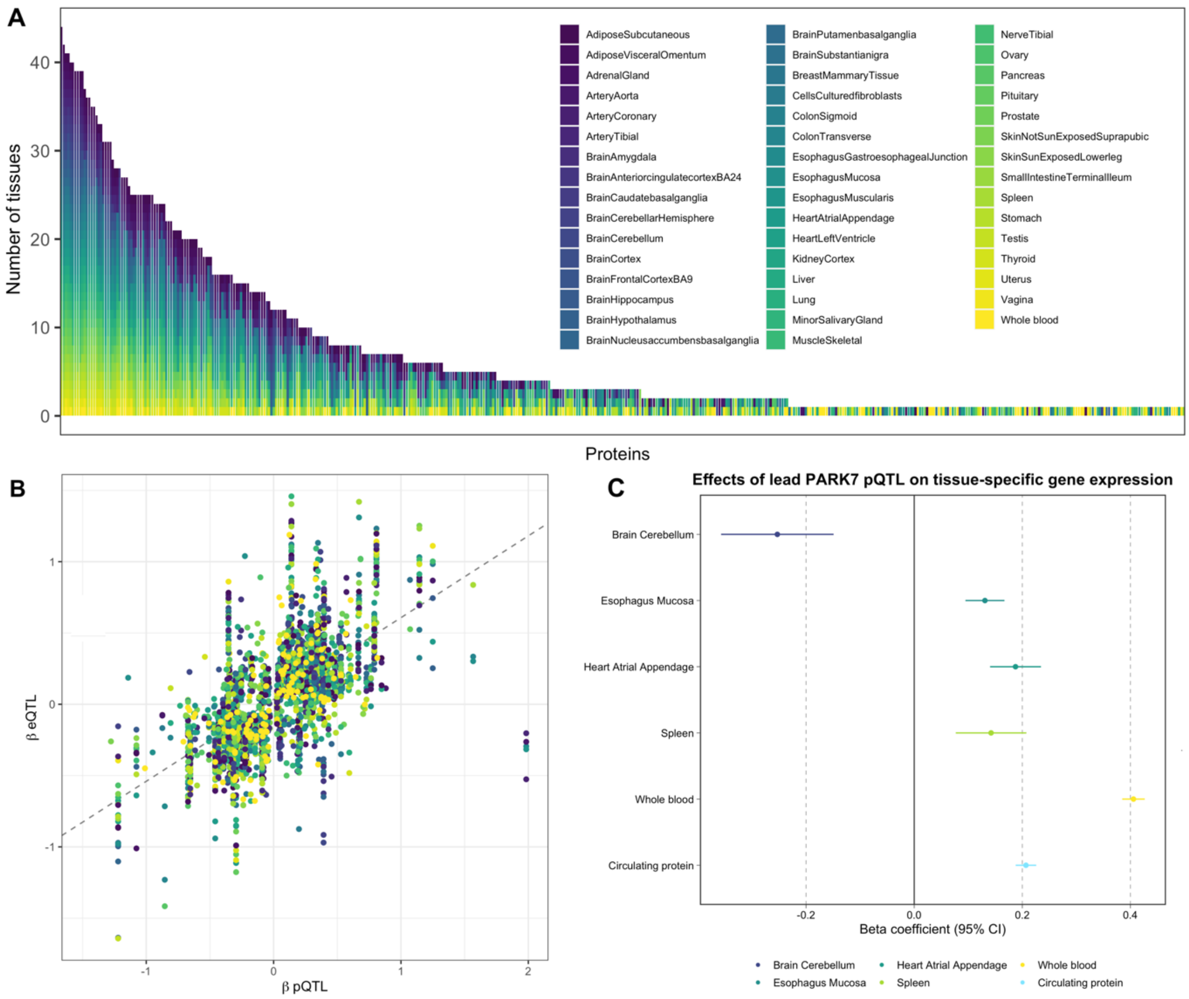

**Extended Data Figure 11. Mendelian randomization estimates of effect of increasing levels of PCSK9 *(cis* genetic instruments) on lipids, cardiovascular disease and stroke risks.**

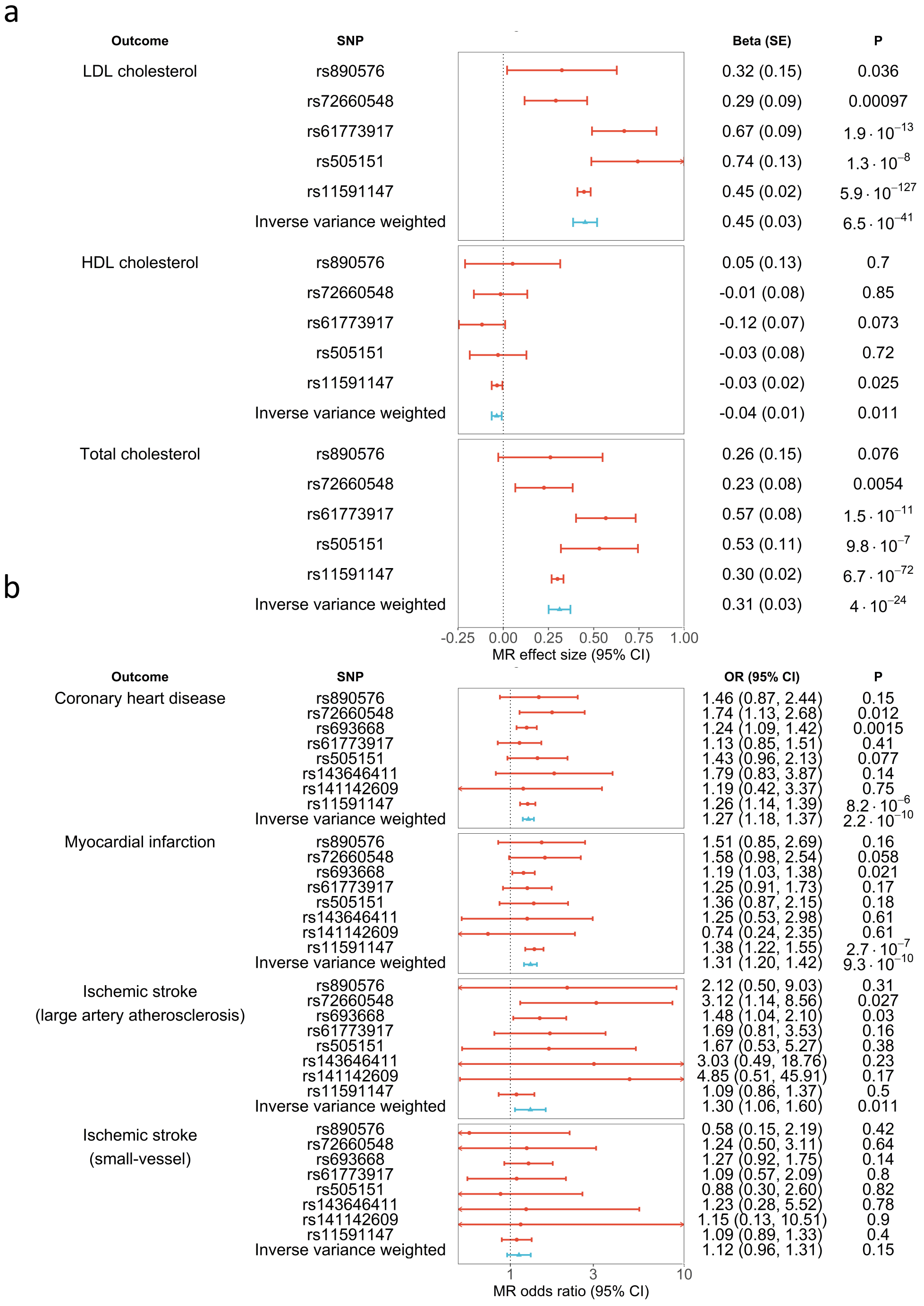
